# Unique mitochondrial carrier has a role in mitochondrial dynamics during Plasmodium falciparum host switching

**DOI:** 10.64898/2026.03.13.711564

**Authors:** Cas Boshoven, Silvia Tassan-Lugrezin, Alannah C. King, Edmund R.S. Kunji, Taco W.A. Kooij

## Abstract

Malaria-causing parasites from the *Plasmodium* genus possess a mitochondrion that is essential across all life-cycle stages and highly divergent from its hosts, making it a suitable drug target. Transport of metabolites across the inner membrane of this metabolically active organelle is mediated by mitochondrial carrier (MC) proteins. Among these, the apicomplexan-specific MC1 (AMC1) stands out due to its likely essential function and limited conservation even within the Apicomplexa phylum. Bioinformatics and structural predictions reveal that *P. falciparum* AMC1 (*Pf*AMC1) lacks canonical gating residues and shows a distorted membrane barrel, suggesting it may not function as a transporter. Using two independent mutant parasite lines, we demonstrate that *Pf*AMC1 localises to the periphery of mitochondria, which exhibit increased dispersion and rounding during male gametogenesis, when the C-terminus is modified. Additionally, we identify the mammalian MTCH2, the function of which is still debated, as a potential structural homologue of *Pf*AMC1, opening new avenues for research. Our findings emphasize the unique role of *Pf*AMC1 in mitochondrial dynamics and lay the groundwork for further exploration of its molecular mechanism.

## Introduction

Malaria is caused by parasites from the *Plasmodium* genus, of which *P. falciparum* (*Pf*) is the deadliest^1^. The number of malaria-related deaths has greatly reduced since the year 2000, but this trend has stagnated in the last ten years and even saw an increase after the COVID-19 pandemic^1^. Additionally, the current rise in resistance against all frontline antimalarials highlights the need for new drugs, preferably targeting new essential pathways and unique proteins to mitigate the risk of cross-resistance^1^.

The mitochondrion is a proven drug target with great potential, as it harbours unexplored unique biology that is different from both its human and mosquito hosts^2–6^. The canonical function of mitochondria is the production of ATP through oxidative phosphorylation (OXPHOS). However, this pathway is only upregulated during the sexual blood stage of the parasite’s life cycle, while the symptomatic asexual blood-stage (ABS) parasites cover their ATP demand mainly through cytoplasmic glycolysis^7^. Nevertheless, the mitochondrion is indispensable during ABS development, as it forms a metabolic hub for a variety of processes, such as the biosynthesis of iron sulphur clusters and ubiquinone, as well as *de novo* pyrimidine biosynthesis^8–11^.

Transport across the permeable outer mitochondrial membrane (OMM) is mediated by relatively large non-selective pores called voltage-dependent anion channels (VDAC) of which one has been identified in *Plasmodium* (PF3D7_1432100)^12^. The critical metabolite exchange over the impermeable inner mitochondrial membrane (IMM) that is required for proper mitochondrial activity is predominantly established by transport proteins belonging to the SLC25 mitochondrial carrier (MC) family^13^. The structure and sequence motifs of MC family proteins are highly conserved, even across very diverse species^14^. MCs are characterized by three sequence repeats of approximately 100 amino acids each. Each repeat consists of two transmembrane α-helices connected by a short amphipathic matrix helix and loop.^15^ The specificity of the substrate binding site is determined by three contact points, one on each even-numbered helix^16–18^. Charged residues of the conserved motifs PX[DE]XX[KR]XXXQ on the odd-numbered helices (matrix side)^15^ and [FY][DE]XX[KR] on the even numbered helices (cytoplasmic side)^16,19,20^ can form salt bridge networks in an alternating manner. These networks are strengthened further by glutamine braces on the matrix side^21^ and tyrosine braces on the cytoplasmic side.^19^ This ensures that the substrate binding site is only ever exposed to one side of the membrane at any time in order to prevent proton leak (**Fig. 1**)^16^. A large number of residues with small side chains facilitate the interconversion between states.^13,19^ Finally, two conserved motifs, [YWF][KR]G and [YF]XG, are involved in the binding of three cardiolipin molecules at the domain interfaces of MC and they are crucial for stability and mechanism of the transporter^15,19,21^.

**Figure 1.**
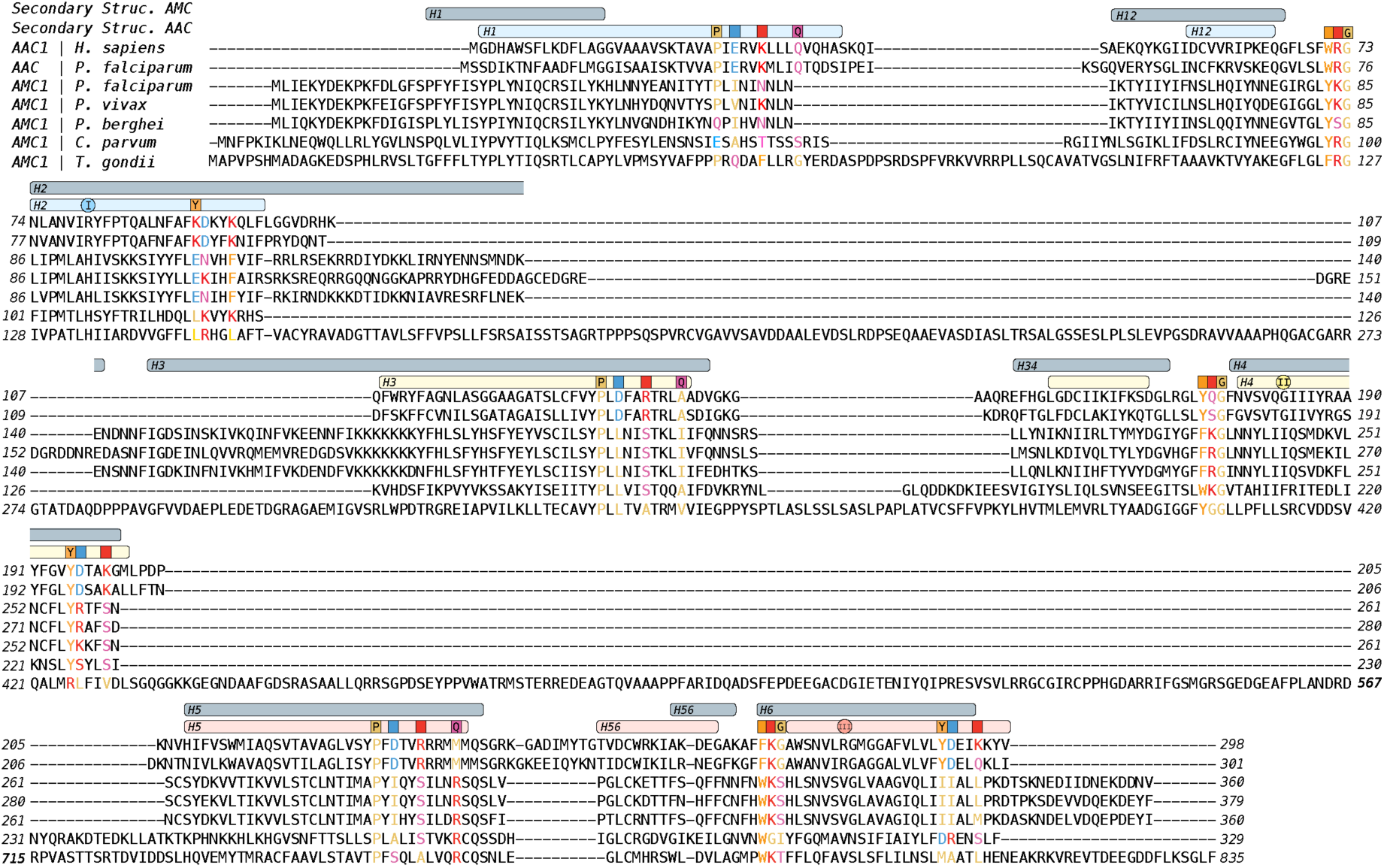
Protein alignment of canonical H. sapiens and P. falciparum (PF3D7_1037300) mitochondrial ADP/ATP carriers (AAC) with P. falciparum (PF3D7_0108800), P. vivax (PVX_081340), P. berghei (PBANKA_0204700), C. parvum (cgd6_2360), and T. gondii (TGME43_276300) AMC1. Whilst there is considerable overlap between the helices of AAC and the predicted helices of AMC1, the latter often lack key residues that are crucial for mitochondrial carrier function. However, there is sufficient sequence conservation between the motifs to suggest that AMC1 is related to the mitochondrial carrier family. The positions of the alpha helices are shown above the alignment for PfAMC1 (grey, predicted using Alphafold) and HsAAC1 (coloured). For HsAAC1, the contact points are marked with a circle and Roman numerals. Key mechanistic and/or structural residues are marked with the canonical amino acid, or with a coloured square representing the biophysical properties of the amino acids canonically found at that position. The amino acids are coloured according to their properties: acidic D, E are blue; basic K, R, H are red; polar C, N, Q, S, T are pink; aliphatic P, G, A, I, L, M, V are yellow; aromatic F, Y, W are orange.

With only 13 members, this family is strongly reduced in *Plasmodium* compared to the 58, 53, and 35 members found in *Arabidopsis thaliana*, *Homo sapiens*, and *Saccharomyces cerevisiae*, respectively^22–24^. There are five carriers in *P. falciparum* with clear sequence homology to well-characterised homologues in model organisms: two ADP/ATP carriers, a phosphate carrier, a dicarboxylate-tricarboxylate carrier, and an S-adenosylmethionine carrier^22^. For these five MCs, experimental evidence has confirmed their predicted metabolite specificities^23,24^, leaving the substrates of the other eight MCs to be elucidated. One member in particular, the *P. falciparum* apicomplexan-specific mitochondrial carrier 1 (*Pf*AMC1), showed extremely limited sequence homology outside of the apicomplexan phylum, giving no lead on the potential function or metabolite transported^25^. Additionally, whole genome essentiality screens in both *P. falciparum* and the rodent malaria model parasite *Plasmodium berghei* indicated an essential function during ABS development^26,27^. This marks *Pf*AMC1 as a unique MC of unknown function, thus meeting the requirements for a potential drug target with limited side effects in humans.

In-house complexome profiling experiments suggested that *Pf*AMC1 may form a complex together with the mitochondrial phosphate carrier (PF3D7_1202200) and two proteins of unknown function, PF3D7_1024500 and PF3D7_0801700^28^. PF3D7_1024500, which is predicted to be mitochondrial (PlasMitoCarta score 2.8, rank 213), shows homology to two mitochondrial proteins with intriguing functions in yeast, Tim21 and Coa1^29^. Tim21 is thought to bring the inner and outer mitochondrial translocases in close proximity and link Tim23 to cytochrome *bc*1-cytochrome *c* oxidase (bc1-COX) supercomplexes^30^. Coa1 is required for cytochrome oxidase complex IV assembly, hinting towards roles in OXPHOS assembly^31,32^. The essentiality and lack of homology of *Pf*AMC1, in combination with the potential new mitochondrial complex drove us to further unravel its function.

Here, we use computational approaches to show that *Pf*AMC1 lacks the structural motifs that are required for a MC-like transport mechanism. By combining experimental genetics and 3D-imaging, we show that *Pf*AMC1 is a mitochondrial protein that localises around fragmented mitochondria in activated microgametocytes, with a potential dual-localisation at the distal end of microgametes. Finally, using novel bioinformatic tools we identified *Pf*AMC1 as a structural homologue of the orphaned MC protein *Mm*MTCH2 and show that *Pf*AMC1 plays a role in mitochondrial dynamics upon male gametogenesis.

## Results

### *Pf*AMC1 is not a canonical mitochondrial carrier family member

Canonical members of the MC family contain three ∼100 amino-acid repeats of the conserved MC domain. The domain prediction tools SMART and InterProScan only predict one MC domain for the 360 amino acids long *Pf*AMC1^33–35^. Reciprocal BLAST analysis of all malaria parasite transport proteins indicated that AMC1 is specific to apicomplexan parasites^25^. Alignment of the AMC1 protein sequences of five Apicomplexa showed a sequence identity of ∼70-75% between different *Plasmodium spp.* but relatively low conservation with the more distantly related *Toxoplasma gondii* (17%) and *Cryptosporidium parvum* (27%) (**Supp. table 3**). Alignment of *Pf*AMC1 with the canonical and best-studied MC family member, the human ATP/ADP carrier (AAC) showed an even lower sequence identity (11%) (**Supp. table 3**). Indeed, at the sequence level, there are very few commonalities between AAC and AMC1 (**Fig. 1**). All three PX[DE]XX[KR]XXXQ motifs were altered with often only the proline retained. A similar level of disruption was seen in the three [FY][DE]XX[KR] motifs, suggesting together that AMC1 would have either very weak or potentially non-functional networks. Interestingly, greater conservation was seen across the cardiolipin binding sites [YWF][KR]G and [YF]XG, suggesting that AMC1 might still be able to bind cardiolipin molecules. With regards to substrate specificity, we saw that contact point I^18,36^ was a conserved positively charged histidine and contact point III was a conserved valine across all AMC1 orthologues. However, contact point II was slightly more variable, generally being a hydrophobic residue^18,36^. Additionally, we compared the structure of the ADP/ATP carrier (AAC) to the AlphaFold model of *Pf*AMC1 to investigate differences at the predicted secondary and tertiary structure level, for which *Pf*AMC1 is predicted to have six transmembrane alpha helices around a central translocation pore^37^. The helices do appear to be of different length as compared to AAC. For example, transmembrane helix H1 was predicted to be shorter in *Pf*AMC1, whilst transmembrane helices H2 and H3 were much longer. *Pf*AMC1 transmembrane helices H5 and H6 were similar in length to those seen in AAC. Taken together, the differences seen in the predicted secondary structure and in the motifs that are known to be required for transport function, strongly suggest that AMC1 is no longer capable of substrate transport. However, the remnants of the motifs and the cardiolipin binding sites show its relationship to the SLC25 mitochondrial carrier family.

### Two independent lines confirm *Pf*AMC1 mitochondrial localisation

Since *Pf*AMC1 lacks the residues and motif required to perform a canonical MC transport function, this raises questions with regards to its molecular function. We were unable to obtain parasites after three independent attempts to delete *PfAMC1*, which is in line with essentiality screens in *P*. *berghei* and *P*. *falciparum*^26,27^. Next, we tagged *PfAMC1* with a triple HA epitope tag along with a glmS ribozyme that allows functional studies using anti-HA epitope antibodies and the induction of a conditional knockdown using the glmS system^38^. We generated two independent lines with slightly different designs (**Fig. 2A&B**). Line 1 (A1.1) harbours a mitochondrial marker in the form of mitochondrial targeted mScarlet (mito-mScar) to enable co-localisation studies^39^. In line 2 (A1.2), we aimed to select for integration of the repair plasmid by replacing the mito-mScar cassette with a human dihydrofolate reductase (h*DHFR*) drug resistance cassette. Therefore, we combined this with a CRISPR-Cas9 guide RNA plasmid with a yeast Dihydroorotate dehydrogenase (y*DHODH*) cassette replacing the h*DHFR* drug resistance cassette (**Fig. 2A**). In the design of line A1.1, we aimed to increase CRISPR-Cas9 efficiency using two different guide target regions. With the few suitable guides available in this region, the second guide was located further in the 3’UTR compared to A1.2, in which only one guide was used (**Supp. fig. 1**). To assess *Pf*AMC1 localisation, we Percoll/sorbitol synchronised parasites in a five-hour window, then took samples for either IFA or SDS-PAGE western blot at ∼10h (ring, R), 24h (trophozoite, T), and 35h (schizont, S) post invasion (**Fig. 2C&D**). Using HA-antibodies, a clear band was observed for both lines at ∼47 kDa (predicted mass AMC1-3xHA = 45.8 kDa), which was absent in the MitoRed control line that harbours a mitochondrial targeted red fluorescent protein mScarlet (**Fig. 2C**)^39^. The signal was strongest in schizonts, while no band was observed in trophozoites and only a faint band in ring stages. Unexpectedly, a much more intense band was observed at ∼160 kDa in line A1.2, which was absent in line A1.1 and the control (**Fig. 2C & Supp. fig. 2**). This unexpected deviation between line A1.1 and A1.2 was also found on IFA. The HA signal in line A1.1 showed a clear co-localisation with mitochondria following the stage specificity as observed in western blot analysis (**Fig. 2D, A1.1**). In line A1.2, however, we found an intense HA signal co-localising with DAPI in all stages (**Fig. 2D, A1.2**). In rings and schizont, we also observed a faint HA signal in DNA-free regions, co-localising with MitoTracker^TM^ Orange (**Fig. 2D, A1.2. zoom-ins**). Upon careful examination of the plasmids used to generate these lines, we realised that Cas9 is N-terminally tagged with a 3xHA. As Cas9 is a nuclear protein of ∼160 kDa, we deem it very likely that the additional HA signal is caused by the presence of 3xHA::Cas9^40^. Western blot analysis stained with three different Cas9 antibodies indeed revealed the higher molecular weight bands to originate from Cas9 in line A1.2 (**Supp. fig. 3A**). Absence of secondary signals in A1.1 indicate that the Cas9-guide plasmid used to generate these lines was exclusively maintained in line A1.2 and lost in A1. The used plasmids only differ in their respective drug resistance cassette, *i.e.* h*DHFR* in A1.1 versus y*DHODH* in A1.2 (**Fig. 2A**). *Pf*DHODH is a mitochondrial protein and known drug target, and complementation with the cytoplasmic y*DHODH* provides resistance to these compounds^41^. Since *Pf*AMC1 is mitochondrial, we hypothesised that the introduction of yDHODH may give a growth advantage when the gene is modified. To test this hypothesis and further characterise *Pf*AMC1, we assessed the growth rates of both lines. We commenced with a titration growth assay on line A1.1 (**Supp. fig. 4A**). At 5 mM glcN, a known critical glucosamine (glcN) concentration for wild-type (WT) parasites, parasite growth was affected though not detrimentally in the first two cycles and parasites did recover after wash-out^38^. We did not observe obvious morphological defects of the mitochondrion as it formed complex branched structures, which segmented normally after treatment with 5 mM glcN for 72h (**Supp. fig. 4B**). To minimize glcN side effects, we opted to assess parasites growth over a longer period with 2.5 mM glcN for both lines. Using 2.5 mM glcN in ring stages, we achieved 80% knockdown in next cycle schizonts in both lines (**Fig. 2E**). Even though parasite growth of glcN treated A1.1 parasites was affected more compared to both the WT and MitoRed controls, parasites continued to proliferate even after four cycles (**Fig. 2F**). Interestingly, untreated A1.2 parasites generally grew slower than the other 3 lines, while the growth factor in the first cycle was less affected by glcN treatment for line A1.2 than for line A1.1 (**Fig. 2F**). Unfortunately, attempts to transfect line A1.1 with a y*DHODH* drug cassette containing plasmid to explore its potential to rescue the growth phenotype were unsuccessful. These data indicate that *Pf*AMC1 might have a role during ABS, but 80% reduction of the protein did not affect parasite survival. Also, the presence of y*DHODH* cassette might reduce the effect of *Pf*AMC1 knockdown.

**Figure 2.**
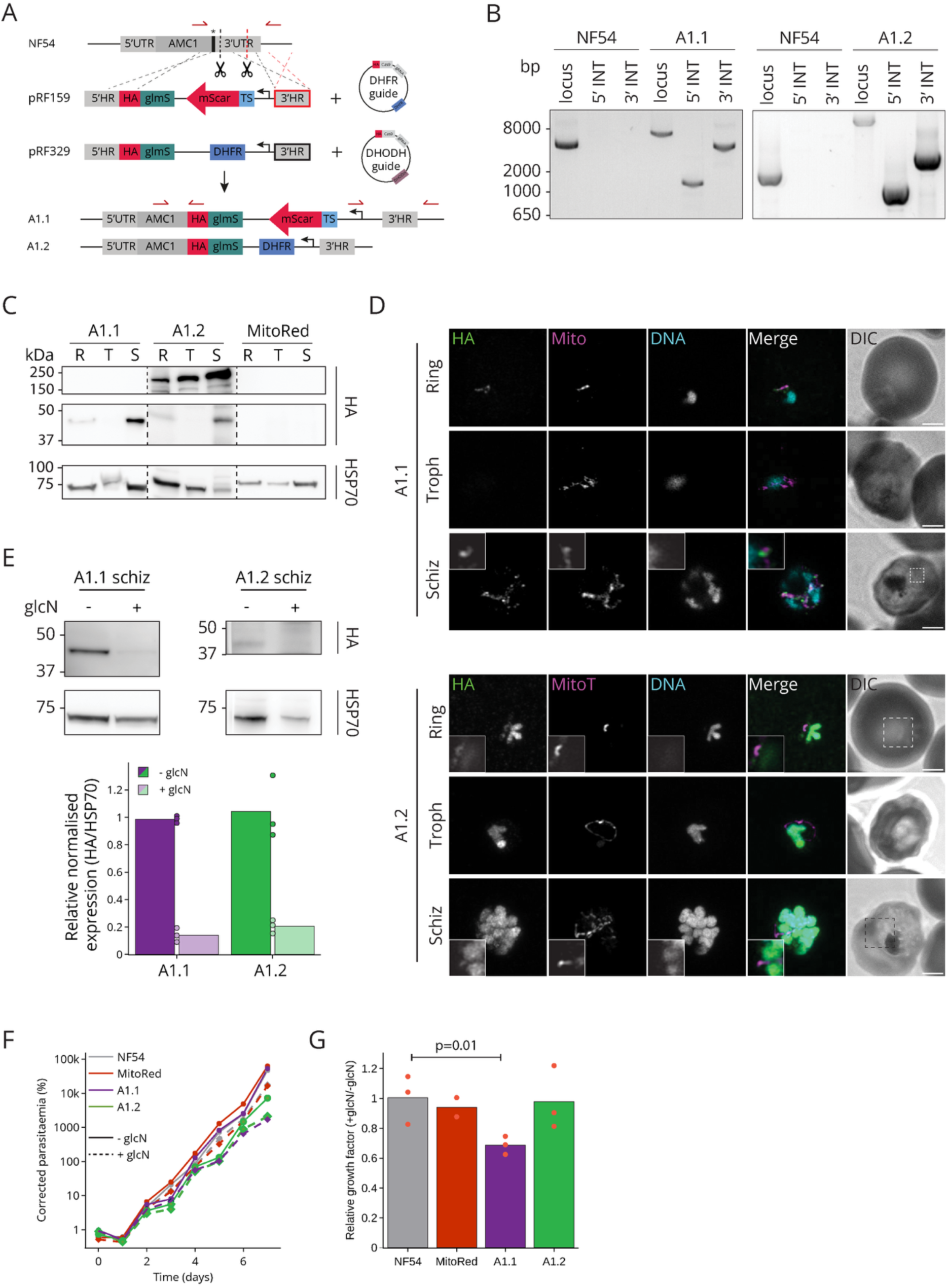
80% reduction of mitochondrial PfAMC1 is not detrimental for parasite growth. A) Schematic overview of repair and guide plasmids used to generate A1.1 and A1.2. The guide RNA and corresponding homology regions (HR) indicated in red for A1.1 and in black for A1.2. B) Diagnostic PCR of lines A1.1 and A1.2 with integration specific primer combinations (indicated in panel A), demonstrating successful 5’ and 3’ integration and the absence wild type (NF54). C) Western blot analysis of saponin lysed ring (R), trophozoite (T), and schizont (S) samples of lines A1.1, A1.2, and a non-HA control (MitoRed). Blots were stained with anti-HA and anti-PfHSP70 antibodies. D) Immunofluorescent assays of ring, trophozoite (troph), and schizont (schiz) samples of lines A1.1 and A1.2. Images are maximum intensity projections of Z-stacks (0.13μm intervals). Both lines were stained with anti-HA (green) and DAPI (DNA, cyan). The mitochondrial signal (mito, magenta) in line A1.1 was captured from the remaining mito-mScar signal. Line A1.2 was stained with MitoTracker^TM^ Orange (MitoT, magenta). Scale bars, 2 μm. Zoom-ins are single slices, depicted by the dashed squares. E) Western blot analysis to determine knockdown efficiency of the 47 kDa band. Parasites were treated with 2.5 mM glcN at ring stage and harvested 70 hours later as schizonts. Average HA intensity was reduced 86% for A1.1 and 80% for A1.2. F) Independent growth assay comparing the growth rate of 2.5 mM glcN treated parasites to a glcN-free control culture. Parasite growth of line A1.1 (purple) and A1.2 (green) was compared to WT (grey) and MitoRed (red). Parasites were Percoll-sorbitol synchronised in a five-hour window and directly set up at ∼0.5% parasitaemia. Samples were taken every 24 hours for seven days. Parasitaemia was determined by flow cytometry and corrected for the dilution factor. G) Effect of 2.5 mM glcN treatment on the growth factor after the first cycle, from three independent growth assays. The effect size is determined by dividing the growth factor (parasitaemia on day 3 / parasitaemia on day 1) of glcN treated samples over the non-treated samples.

### Knockdown of *Pf*AMC1 does not affect sensitivity to OXPHOS targeted drugs

Three out of four proteins in the putative *Pf*AMC1 complex are uncharacterised. With few leads on the potential function of this complex, it is interesting that PF3D7_1024500 shows domain similarity to Tim21 and a cytochrome oxidase complex IV assembly protein COA1^28^, both of which have a role in the assembly of OXPHOS complexes^30–32^. We hypothesised that if *Pf*AMC1 is indeed in a complex together with OXPHOS complex assembly proteins, knockdown of *Pf*AMC1 could make parasites more susceptible to OXPHOS targeting drugs but leave them similarly susceptible to non-OXPHOS targeting drugs, as shown previously^42,43^. To test this, we treated A1.1 parasites with 2.5 mM glcN to induce *PfAMC1* knockdown and compared the drug-sensitivity to untreated A1.1 parasites and WT controls with and without glcN treatment. We included DSM-1 and DSM265, targeting *Pf*DHODH, atovaquone, ELQ-300 and proguanil, targeting complex III, and chloroquine, dihydroartemisinine, and MMV693183, targeting non-mitochondrial pathways (**Fig. 3**)^44–47^. Since A1.2 was found to harbour a y*DHODH* resistance cassette, which potentially interfered with the read out, we excluded this line from the assay (**Supp. fig. 4C**). To our surprise, none of the compounds caused a shift in EC_50_ upon glcN treatment of A1.1 (**Fig. 3**). Even though some shifts were significant, all effect sizes were below one order of magnitude and, therefore, regarded as irrelevant in this biological context (**Fig. 3**). These data provide no clear indication for the involvement of *Pf*AMC1 in the assembly or functioning of OXPHOS complexes nor that parasite health is otherwise affected in such a way that the parasite becomes more sensitive to any drugs.

**Figure 3.**
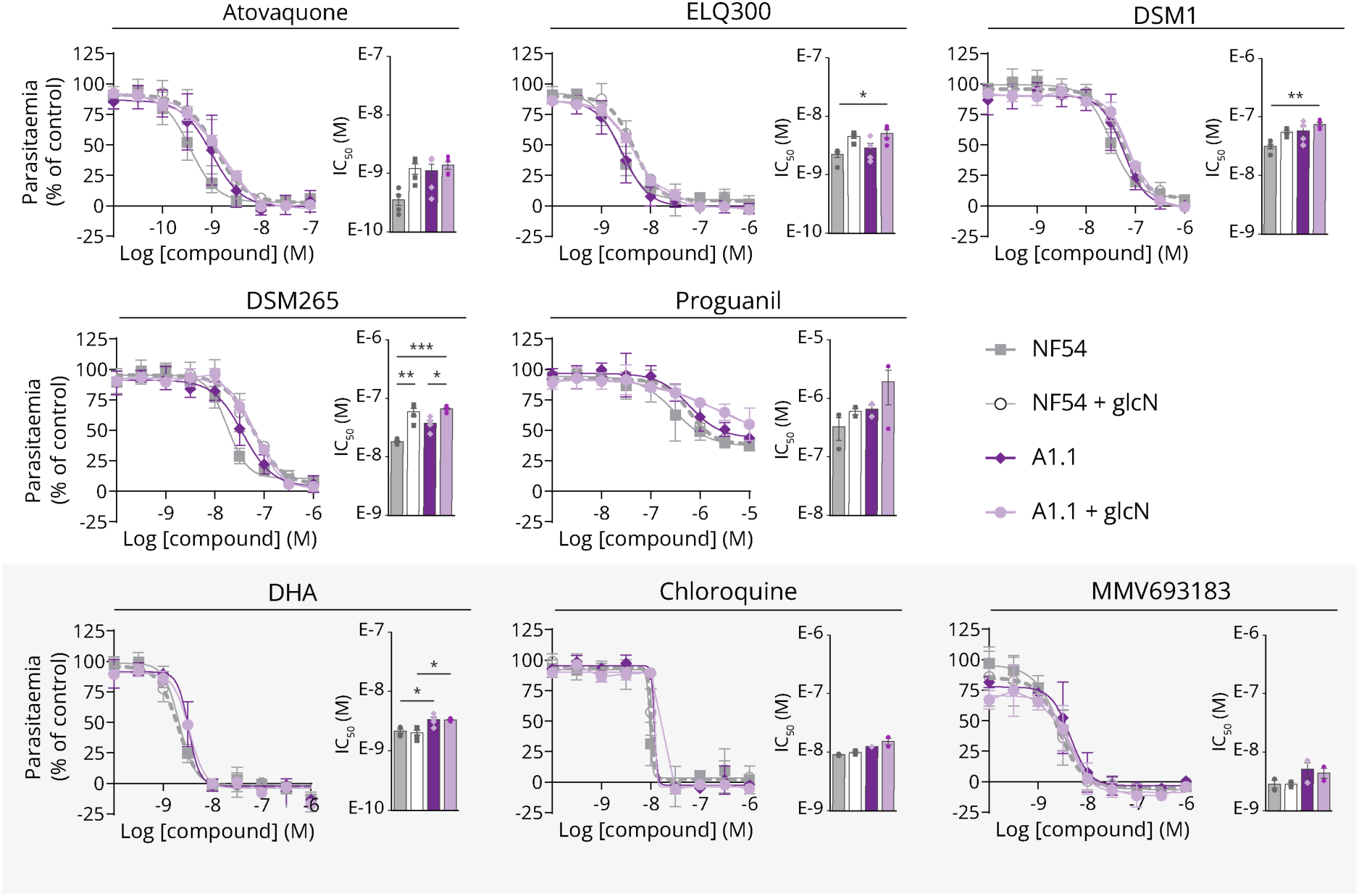
Glucosamine treatment of A1.1 does not affect drug sensitivity. Drug sensitivity profiles and IC50 values for WT and A1.1 mixed ABS parasites, either untreated or treated with 2.5 mM glcN (+ glcN). Profiles for drugs with mitochondrial targets are depicted in a white background, other drugs are shown in a light grey background. The depicted values represent the average value for mean parasite density relative to controls ± SEM. Proguanil, chloroquine, and MMV693183 were tested twice, all other compounds were tested in four independent experiments.

### *Pf*AMC1 localises around fragmented mitochondria in activated microgametocytes

As our initial protein characterisation and drug sensitivity assays in ABS did not yield major insights into the functioning of *Pf*AMC1, we explored a possible role during sexual-stage development. Both A1.1 and A1.2 produced healthy looking stage V gametocytes, in which *Pf*AMC1 localised to the mitochondrion as we observed in ABS (**Fig. 4A&D**). The nuclear Cas9-derived signal observed in A1.2 ABS was still present in stage V gametocytes, however, less intense. Reassuringly, SDS-PAGE western blot analysis of A1.2 showed that the 160 kDa band, corresponding to Cas9, is more abundant in ABS compared to stage V gametocytes or xanthurenic acid (XA) activated gametes (**Supp. fig. 2**). Interestingly, A1.1 stage V gametocytes were unable to exflagellate upon activation with 50 μM XA for 15 minutes, regardless of knockdown (**Fig. 4E**). Even though exflagellation levels in line A1.2 were variable, this line did consistently exflagellate (**Fig. 4E**), highlighting another important discrepancy between the two lines. Immunofluorescence imaging of activated A1.1 parasites showed that male gametes rounded up and expanded their DNA, however, no clear axonemal structures were formed (**Fig. 4A&G, Supp. fig. 5**). Sporadically, we observed thick protrusions also known as a superflagella yet never properly shaped or detached microgametes (**Supp. fig. 5**). In the absence of axonemes, we differentiated males and females based on mitochondrial fragmentation for males versus a seemingly connected and condensed structure in females^39,48^. This separation always correlated to a ring-like tubulin signal in the males versus a dispersed tubulin signal throughout the whole cell for the females (**Fig. 4A, Supp. fig. 5A**). While total HA signal intensities were similar in both sexes, they significantly increased in activated gametocytes compared to stage V gametocytes (**Fig. 4A&B**). In activated male gametocytes, *Pf*AMC1 localised consistently around fragmented mitochondria (**Fig. 4A, C, D, F&G, Supp. fig. 5A-C**). 3D reconstructed Z-stack images of line A1.1 further supported this observation of *Pf*AMC1 encapsulating the matrix localised mito-mScar (**Fig. 4G, Supp. fig. 6, Supp Video 1&2**). Even though mitochondrial carriers are generally localised to the IMM, this observation suggests a potential OMM localisation for *Pf*AMC1.

**Figure 4.**
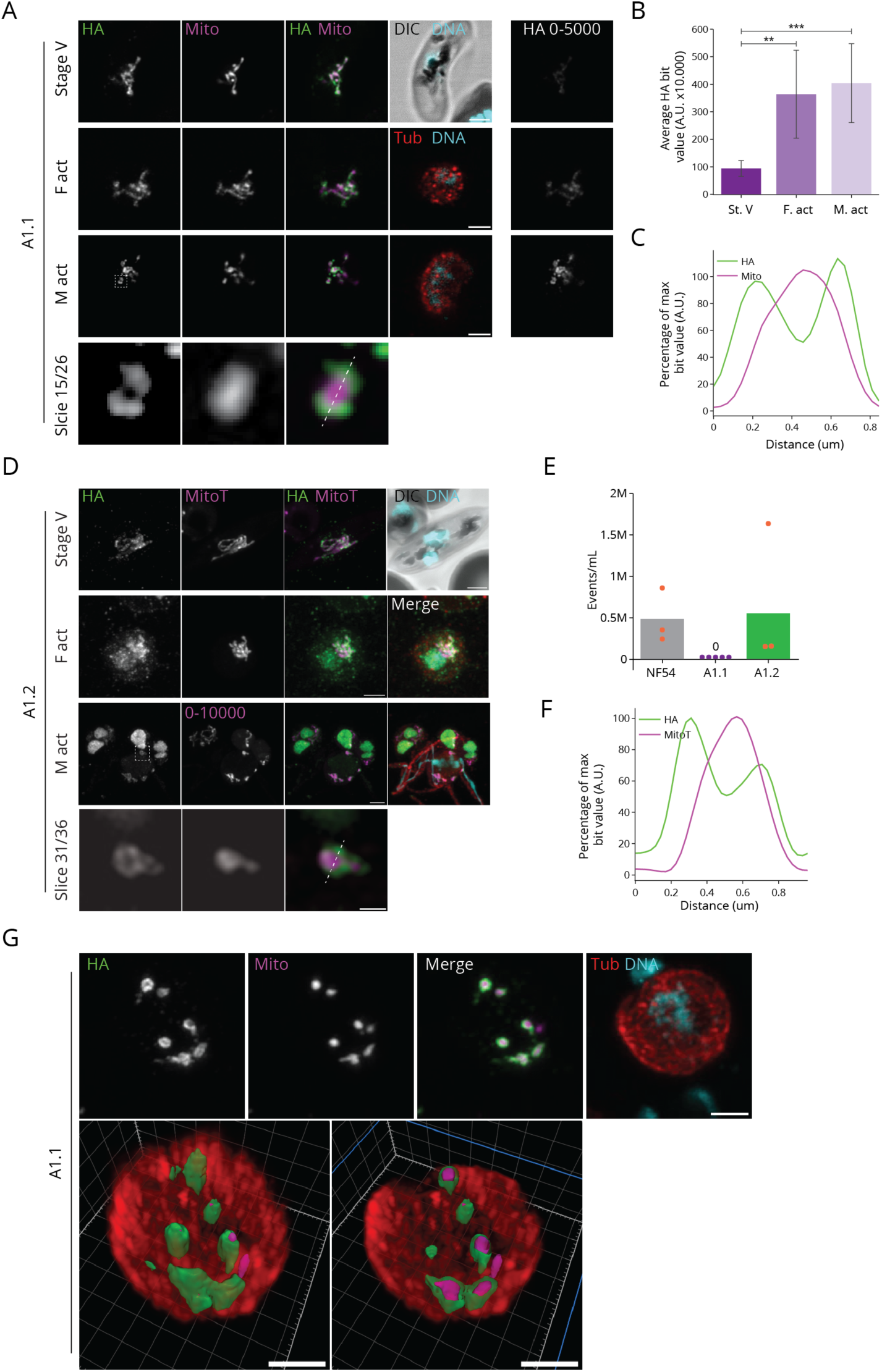
PfAMC1 localises around fragmented mitochondria in activated male gametocytes. A&D) Immunofluorescence imaging of line A1.1 (A) and A1.2 (D) stage V gametocyte and activated female (F) and male (M) gametocytes, stained with anti-HA (green), anti α-tubulin (red) and DAPI (DNA, cyan). A) The mitochondrial signal (mito, magenta) was captured from the remaining mito-mScar signal. Male and female parasites were grouped according to mitochondrion morphology and α-tubulin staining (Supp. fig. 5A). The HA signal histograms were individually adjusted to optimise signal to noise ratios for co-localisation purposes. To compare intensity values, the HA signal is depicted with the same histogram settings (HA 0-5000). Slice 15/26 shows a single slice zoom-in from the activated activate male gametocyte, indicated by the white dashed square. Scale bars, 2 µm. B) HA signal quantification of stage V and activated male and female gametocytes (complete Z-stacks for 10 individual cells per stage). C&F) Signal intensity profiles of HA and mito, plotted over the white dashed line indicated in panels A and D for A1.1 (C) and A1.2 (F), respectively. Values are depicted as percentage of the maximum measure bit value. D) Male and female parasites were grouped according to α-tubulin staining. The mitochondrion was stained with MitoTracker^TM^ Orange (MitoT, magenta) Scale bars, 2 µm. Slice 31/36 shows a single slice zoom-in from the activated male gametocyte, indicated by the white dashed square, scale bar, 0.5 µm. E) Total exflagellation centres for wild type (NF54), A1.1, and A1.2 gametocytes after 15 minutes activation with 50 µM XA, counted on a haemocytometer. G) 3D visualisation of an A1.1 activated male gametocyte, to reconstruct the localisation of PfAMC1 (HA, green) with respect to the matrix localised mito-mScar (Mito, magenta). Scale bars, 2 µm. The numbers behind the channels indicate the histogram settings.

### A tubulin-free compartment at the distal end of microgametes shows HA staining in line A1.2

To our surprise, when imaging exflagellating A1.2 males, we observed a ring-like HA signal around an increased MitoTracker^TM^ signal on one end of free microgametes (**Fig. 5A&B, Supp. fig. 6**). The HA and MitoTracker^TM^ signals seemed to come from a separate structure that was devoid of α-tubulin and DNA. It is important to note that line A1.2 harbours both a nuclear and mitochondrial HA. The MitoTracker^TM^-encircling HA signal is reminiscent of the *Pf*AMC1 localisation at the periphery of the fragmented mitochondrion we observed in A1.1 and A1.2 activated male gametocytes. Additionally, we once observed an A1.1 activated male gametocyte with a single extended axoneme (**Supp. fig. 5A**). Here, one of the mitochondrial fragments seemed to be outside of the residual body, in the same plane and location as where the axoneme appeared to be attached to the residual body. WT microgametes do not appear to harbour a mitochondrion and mtDNA inheritance in *Plasmodium gallinaceum* is maternal^49,50^. Therefore, we wondered whether 3xHA tagging *Pf*AMC1 resulted in a mitochondrial harbouring microgamete potentially due to *Pf*AMC1 malfunctioning. We re-analysed line A1.2 microgametes still attached to the residual body and consistently observed proximal and comparable HA and MitoTracker^TM^ signals at the distal end of microgametes and hence at the opposite side of where the mitochondrial fragment was observed at the base of the attached A1.1 microgamete (**Supp. fig. 5A**). We then imaged MitoTracker^TM^ stained microgametes from WT parasites (**Supp. fig. 6**). This showed that WT parasites also have a structure with increased MitoTracker^TM^ signal at the distal end of the microgamete that is devoid of α-tubulin and partially stains for the cytoplasmic male marker PF3D7_1325200 (**Supp. fig. 6**)^51^. However, in MitoRed parasites, in which the mitochondria are visualised by a mitochondrial localised mScarlet protein, no mitochondrial signal was observed in the tubulin devoid compartment (**Fig. 5D, Supp. fig. 6**)^39^. Earlier studies showed that the basal body is located at the distal tip and these appear as an α-tubulin devoid region in fluorescent microscopy and can be visualised using the NHS-ester stain in expanded microgametes^49,52^. Expanded MitoRed activated male gametocytes stained with MitoTracker^TM^ and NHS ester indeed revealed the basal bodies (**Fig. 5E**). This showed that the basal body and mitochondria rarely co-localise. In two incidents, these structures appeared to co-localise in maximum projection images but were shown to be separated in the Z-direction (**Fig. 5E, Supp. fig. 6**). Upon adjusting brightness and contrast, we observed a specific MitoTracker^TM^ signal at the basal body. A similar signal was not found in non-expanded MitoRed activated male gametocytes in which the mitochondria were visualised by mito-mScar (**Supp. fig. 6**). This indicates the MitoTracker^TM^ signal from the expanded cells does not come from mitochondria but potentially indicates a slight membrane potential around the basal body. As our western blot analysis demonstrated the presence of HA-tagged AMC1 and Cas9, we tested two different anti-Cas9 antibodies on A1.2 schizonts. Unfortunately, both resulted in background signals only and did not give the expected nuclear stain (**Supp. fig. 3B**). Therefore, the origin and function of the HA signal around the basal body in line A1.2 microgametes remains to be elucidated.

**Figure 5.**
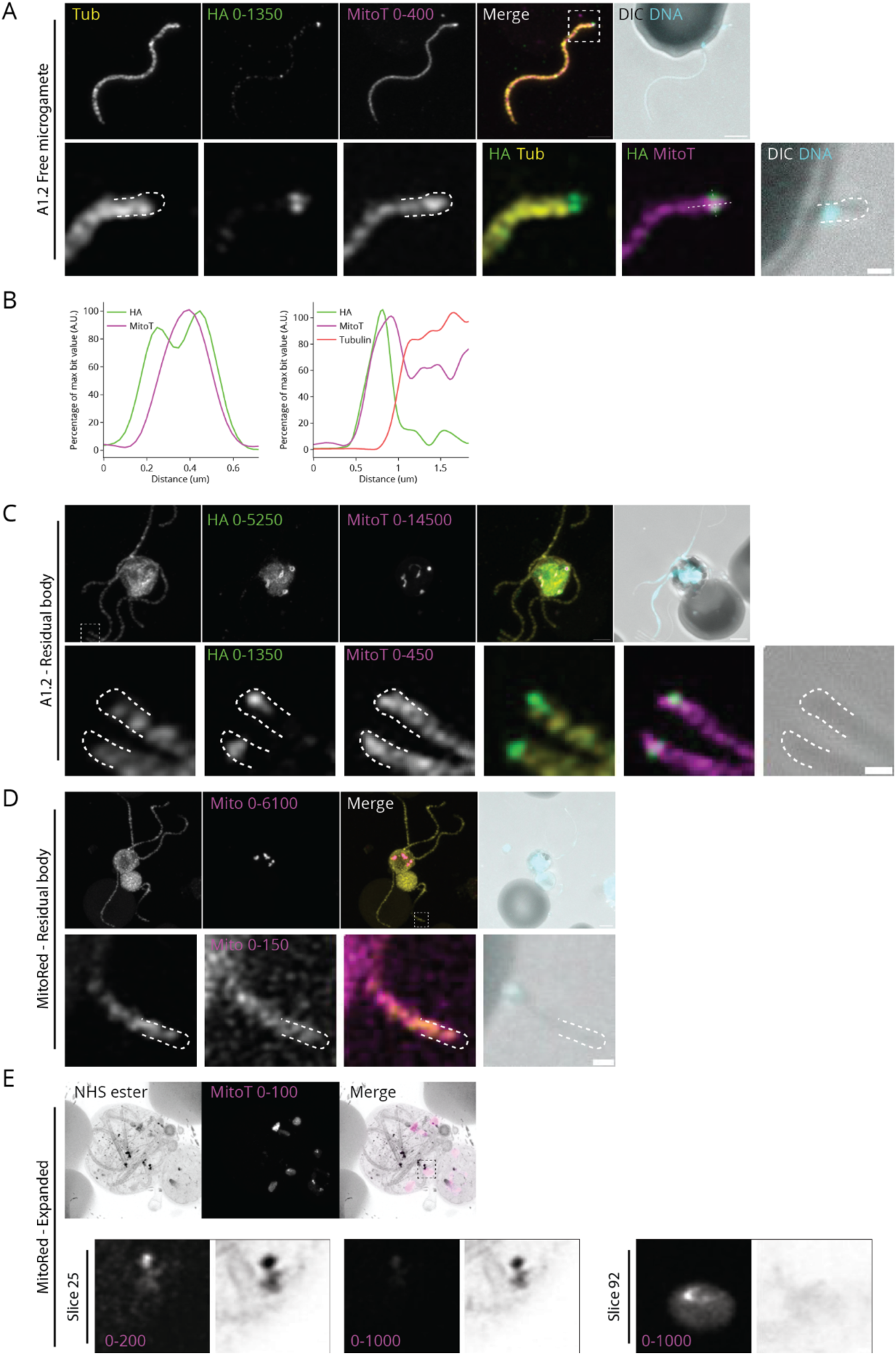
Tubulin-free compartment at the distal end of free microgametes shows HA staining in line A1.2. A, C-E) Maximum intensity of Z-stack image. Scale bar, 2 µm. Zoom-in indicated by the dashed square is a single slice. Scale bars, 0.5 µm. A&C) Line A1.2 free microgamete (total 4/4 microgametes, **Supp. Fig. 6**) and microgametes still attached to the residual body (C) stained for α-tubulin, HA, MitoTracker ^TM^ Orange and DAPI. B) Signal intensity profiles for tubulin (right, horizontal line), HA and mito (right and left, vertical line), plotted over the white dashed lines. D) WT free microgametes (total 7/7 microgametes, **Supp. Fig. 6**) (D) stained for α-tubulin, male marker, MitoTracker Orange, and DAPI. E) Expanded MitoRed activated male gametocytes stained with NHS-Ester and MitoTracker^TM^ Orange. The numbers behind the channels indicate the histogram settings.

### 3xHA tagging *Pf*AMC1 causes mitochondria to round and fragment upon male gametogenesis

We have shown previously that in contrast with ABS that have a single mitochondrion, gametocytes harbour 4-8 tightly connected mitochondria that fragment and round-up during early male gametogenesis^39,48^. As mitochondrial fragmentation was more obvious in both A1.1 and A1.2 microgametocytes and the individual mitochondria appeared rounder compared to WT, we further quantified these observations using the Mitochondria-Analyzer plugin in Fiji^53^. This confirmed that both lines had more individually recognisable mitochondria that are both smaller and more spherical compared to the WT control (**Fig. 6A&B**). As described earlier, the design of A1.1 and A1.2 lines differs in several places. However, both lines show a similar aberrant mitochondrial morphology upon male gametogenesis independently of knockdown. This suggest that the 3xHA tag on the C-terminus of AMC1, the only shared feature between the two lines, is responsible for this phenotype, likely explained by an impaired function of *Pf*AMC1.

**Figure 6.**
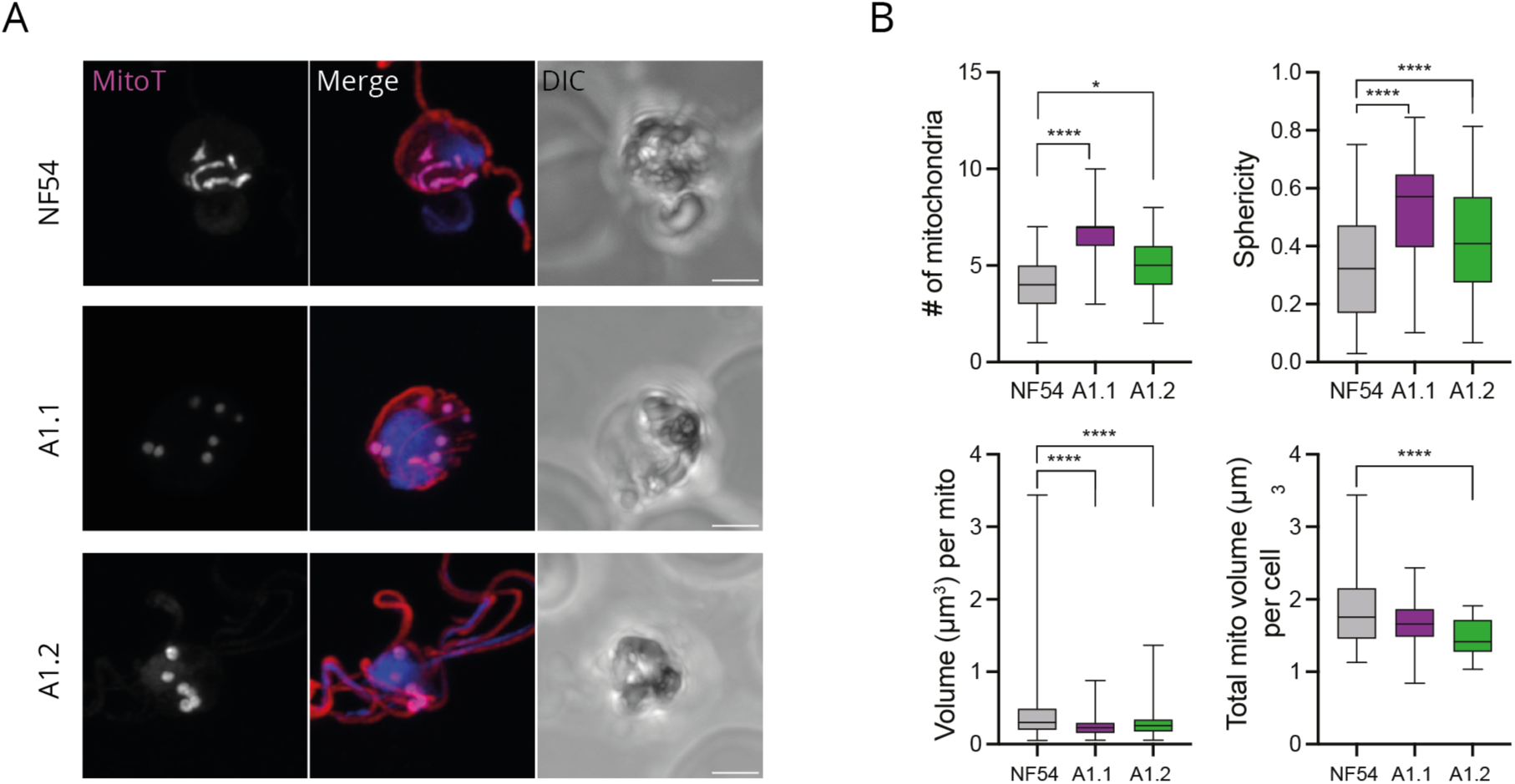
Increased mitochondrial rounding and fragmentation in 3xHA tagged PfAMC1 activated male gametocytes. A) Representative immunofluorescence images of WT, A1.1, and A1.2 microgametes after 20 minutes. XA activation. All samples are stained with MitoTracker^TM^ (magenta), anti-α-tubulin (red) and DAPI (blue). Scale bar, 2 µm. B) Quantification of the mitochondrial morphology using the Mitochondria-Analyzer plugin in Fiji. Z-stack images from two replicates were analysed (WT; N=44, A1.1; N=39, A1.2; N=40). Statistical significance was evaluated by one-way ANOVA followed by Šidák correction. * for P ≤ 0.05, **** for P ≤ 0.0001.

### *Pf*AMC1 is a potential structural homologue of MTCH2

Even though we found the novel and unique localisation for *Pf*AMC1 around more fragmented and rounder mitochondria its function remained elusive. Therefore, we employed recently developed bioinformatic tools to further clarify its function. We used the predicted AlphaFold 3.0 structure as input for the Foldseek protein alignment tool^54,55^. Foldseek is an alignment tool that compares tertiary structures of the input protein to a database of predicted and known structures. This approach yielded the *Homo sapiens, Pongo abelli, Bovis taurus* and *Mus musculus* MTCH2 proteins as potential structural homologues to *Pf*AMC1, which were not identified through protein sequence BLAST (**Table 1**). Protein sequence alignment indeed showed very limited homology to the Foldseek identified structural homologues, aside from the potential cardiolipin binding sites and a few conserved residues in the carrier domain similar to the human AAC discussed above (**Fig. 1, Fig. 7A**). *Pf*AMC1 and *Mm*MTCH2 align well (root mean square deviation (RMSD) *Pf*AMC1 vs *Mm*MTCH2 = 3.9 Å, *Pf*AMC1 vs *Sc*AAC2 (PDB 4C9G) = 4.9 Å), particularly with regards to the unconventional groove, highlighted by the yellow lines in the electrostatic overview, which sets them apart from other members of the mitochondrial carrier family (**Fig. 7B&C**). This is particularly intriguing as these are AlphaFold predictions based on non-homologous protein sequences (**Fig. 7A**). However, where the groove in MTCH2 is created by the lack of transmembrane helix H6, in *Pf*AMC1 it is due to helix H6 angling outward, instead of closing the barrel. The groove is predicted to have a functional role in the ascribed lipid flipping and protein insertion functions of MTCH2^56–58^. While the predicted structures for *T. gondii* (TGME49_276900) and *C. parvum* (cgd6_2360) homologues of AMC1 are far more distinct, these also show a disrupted barrel with a conserved groove in the former and a break in two of the transmembrane helixes in the latter (**Supp. fig. 7**). MTCH2 is an orphaned MC that is localised to the OMM^59^. Two independent groups have shown that deletion of *MTCH2* caused mitochondrial fragmentation and rounding in mouse embryonic fibroblasts and human HCT cells, comparable to what we find upon male gametogenesis of 3xHA tagged AMC1^60,61^ hinting at possible functional homology.

**Figure 7.**
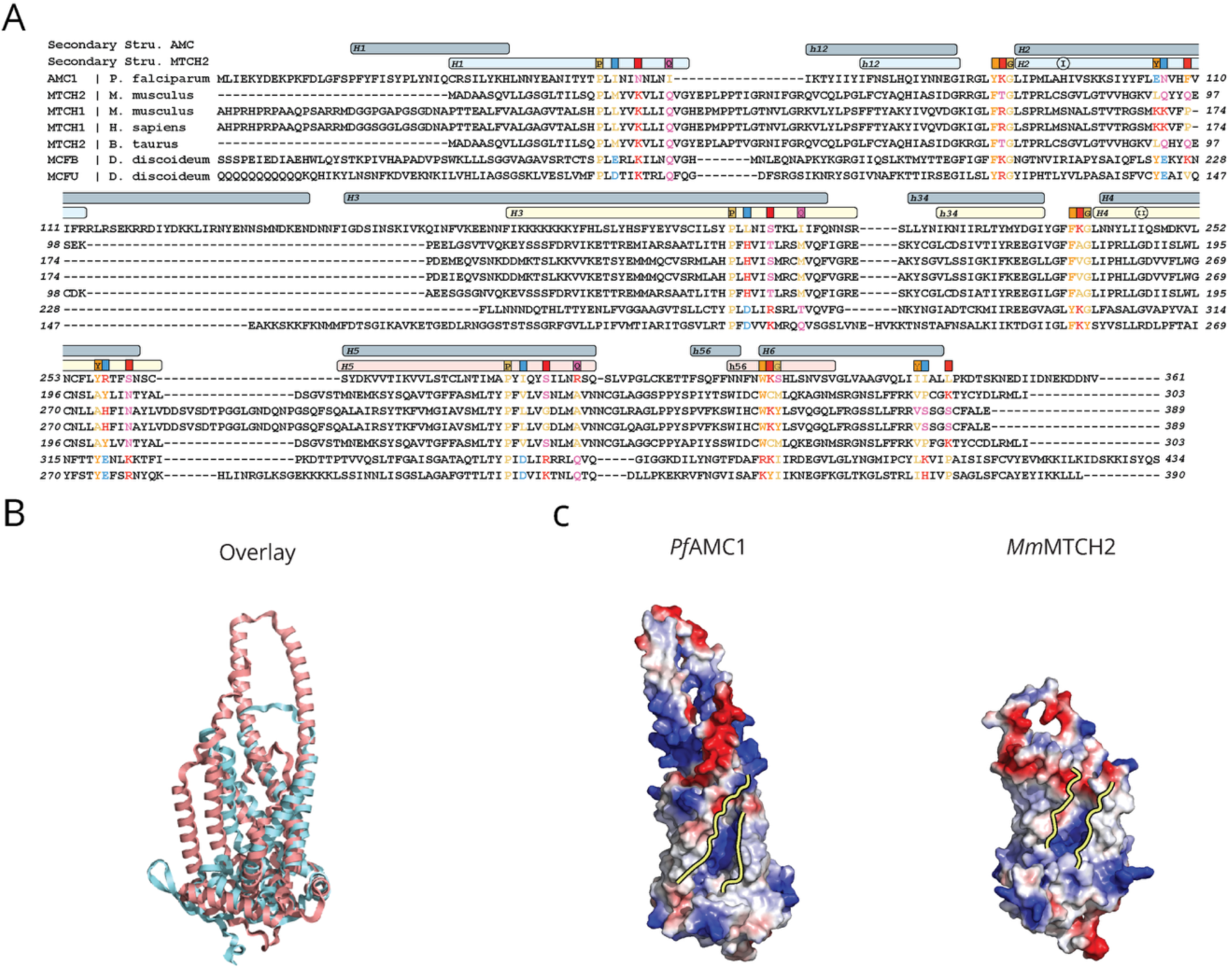
Structural homology between PfAMC1 and MmMTCH2. A) Protein sequence alignment of the Foldseek top hits against the predicted PfAMC1 structure as input. B) Overlay of the predicted PfAMC1 (red) and MmMTCH2 (blue) structure (RMSD = 3.9 Å). C) Electrostatic overview of the predicted PfAMC1 and MmMTCH2 structure with the groove highlighted by the yellow lines.

**Table 1.**
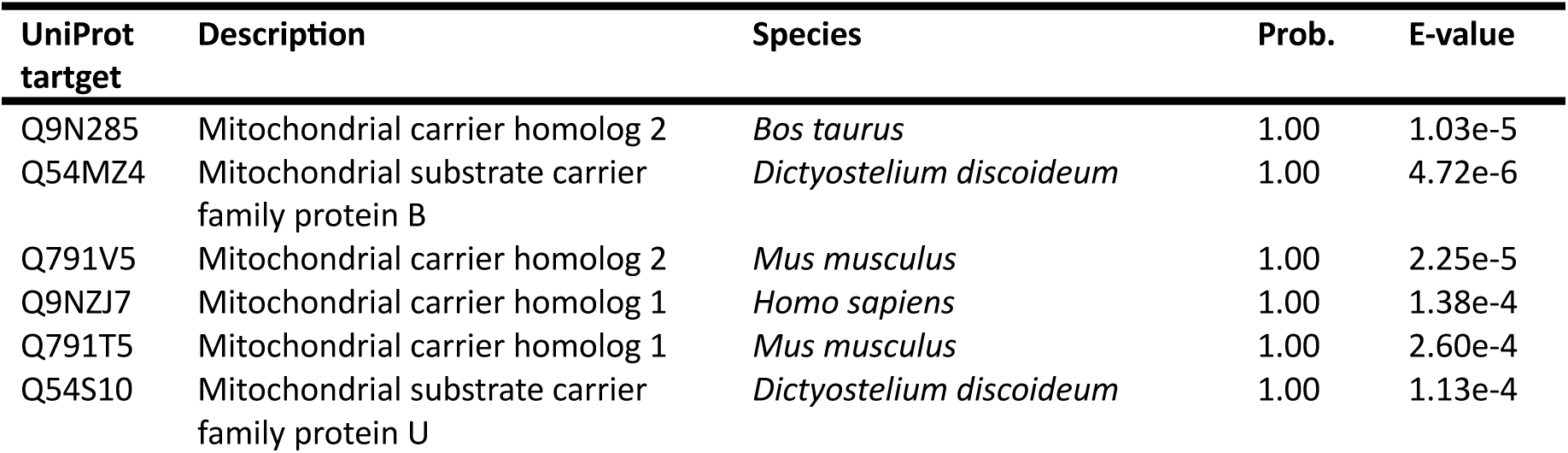
Summary of Foldseek hits against the Swissprot database using AlphaFold 3.0 *Pf*AMC1 (PF3D7_0108800, Q8I254) structure predictions as input.

## Discussion

Mitochondrial carrier proteins are responsible for the transport of metabolites across the impermeable IMM. There are only 13 MC family members in *Plasmodium* forming a ‘minimally viable’ set compared to the 58, 53, and 35 members in *Arabidopsis*, human and yeast, respectively^62,63^. It is therefore intriguing to find that *Pf*AMC1, studied here, lacks the amino acid residues required to perform a MC transport mechanism (**Fig. 1**)^64^. This brings the number of potential transporters in the *Plasmodium* MC family down to 12. The likely essentiality of *Pf*AMC1 in combination with the limited conservation, even within the apicomplexan phylum, highlights the importance of identifying its molecular function.

With the difference in growth defect upon *Pf*AMC1 knockdown and the differing ability of stage V gametocytes to exflagellate, the presented data reveal phenotypic differences between lines A1.1 and A1.2 (**Fig. 2F & 4E**). We identified two major differences in the genetic design of the two lines that might explain the affected phenotypes; the additional deletion of the 3’UTR in line A1.1 compared to A1.2 (Pf3D7_01_v3: 360774..361580, **Supp. fig.1**) and the presence of the y*DHODH* resistance cassette on the Cas9-guide plasmid used to generate line A1.2.

First, the Pf3D7_01_v3: 360774..361580 locus is particularly interesting as it does not cover any annotated genome fragments. However, RNA sequencing revealed male gametocyte-specific transcription activity on the negative strand^65,66^. To assess whether this locus indeed plays a vital role in microgamete formation one could selectively delete this non-coding region and quantify exflagellation in the resulting parasite line. The gene downstream of *Pf*AMC1, *Pf*PHIL1, is a well-characterised IMC protein that is shown to have an essential role in gametocyte development^67–69^. Potential transcriptional fluctuations due to modification in the promoter region could explain the reduction in α-tubulin formation and the lack of exflagellation in line A1.1. However, the deleted 3’UTR fragment in line A1.1 is 1.8 kb away from the *Pf*PHIL1 coding region and only marginally affects the first sex-specific positive strands transcription profiles. Also, the *Pf*PHIL1 ABS transcription profiles only start 1.6 kb after the deleted fragment. Therefore, we consider it unlikely that the phenotypes are caused by downstream effects on *Pf*PHIL1.

Western blot analysis showed the presence of Cas9 in line A1.2, which is transcribed from the CRISPR-Cas9 guide plasmid. An explanation for maintaining the guide in line A1.2 but not in A1.1 could be the y*DHODH* resistance cassette. In the cytoplasm, yDHODH produces orotate, a precursor for pyrimidine biosynthesis^41^. This functionally bypasses the mitochondrially localised *Pf*DHODH, which is the only essential step of the OXPHOS in ABS parasites. It thereby confers resistance to DSM265 and atovaquone, targeting *Pf*DHODH and the cytochrome *bc*_1_ complex, respectively^5,41^. The absence of phenotypes in the y*DHODH* harbouring line A1.2 make it tempting to speculate that introducing y*DHODH* rescued the phenotypes observed line A1.1. This could hint towards a primary or secondary role of *Pf*AMC1 in pyrimidine biosynthesis. To test this hypothesis, we attempted to transfect two different y*DHODH* harbouring plasmids into the A1.1 background, hypothesising this would rescue the growth and exflagellation phenotype. Unfortunately, three transfection attempts did not yield any viable parasites.

Line A1.1 does also harbour a mitochondrially localised mScarlet. However, as the MitoRed line, which harbours the same cassette integrated in a silent locus, does not show aberrant growth compared to WT upon 2.5 mM glcN treatment and demonstrates normal ABS and sexual blood-stage development, this cassette is unlikely to contribute to the observed phenotype (**Fig. 1E**)^39^.

### HA staining on the distal tip of microgametes

An exciting observation is the HA signal at the distal tip of A1.2 microgametes (**Fig. 5A**). With this comes the important question on the origin of the signal as these parasites showed both a nuclear and a mitochondrial fraction. One possibility is the nuclear Cas9, which seems unlikely as no HA signal is observed in the microgamete nucleus and the *Plasmodium* basal body is assembled in the cytoplasm (**Fig. 5A**)^49,70,71^. The other option is the mitochondrial *Pf*AMC1, which is intriguing to speculate as we observed the HA signal around a weak MitoTracker^TM^ signal in line A1.2 and an extracellular mitochondrion attached to an axoneme in line A1.1. Earlier electron microscopy, genome and proteomic studies showed that *Plasmodium* microgametes do not harbour a mitochondrion^49,50^. The extracellular mitochondrion in line A1.1 initially led us to hypothesise that the C-terminal 3xHA tag on *Pf*AMC1 caused a connection between the axoneme and the mitochondrion (**Fig. 5A, Supp. fig. 6A**). We rejected this hypothesis as we showed that the mitochondrial signals in the two different lines to belong to the opposite ends of the microgametes. We also found WT parasites to harbour a MitoTracker^TM^ signal at the distal tip, with lower signal intensity compared to the mitochondrial fragments in the residual body (**Fig. 5**). Imaging of WT microgametes and expanded MitoRed activated male gametocytes indicated that the MitoTracker^TM^ signal came from the basal body (**Fig. 5**). As MitoTracker^TM^ concentrates in areas with a positive membrane potential, it is intriguing to speculate such a potential is present around the basal body. However, how this is accomplished or relevant remains to be elucidated as the ATP required for the flagellar movement through the dynein motor is thought to be produced through glycolysis, which does not require a membrane potential^72^. Due to the lack of a lactate exporter, Talman *et al*. speculated on the cytoplasmic acidification of microgametes, potentially explaining the MitoTracker^TM^ enrichment^72^. This work shows the dependency of microgamete movement on glycolysis, even though 56/615 identified proteins in the microgamete proteome are predicted mitochondrial^29,72^. Recent work shows that mitochondrial ATP production is also important for male gametogenesis in *P. falciparum*^51^. An alternative explanation for *Pf*AMC1 to be on the distal tip is a dual localisation apart from the mitochondrion specifically in microgametes. For this it is worth noting that the *Pf*AMC1 *P. berghei* homologue (PBANKA_020470) is not identified in the microgamete proteome dataset^72^. To speculate on the potential role of *Pf*AMC1 at the distal tip we would have to understand the function of this protein better.

### *Pf*AMC1 as structural homologue to MTCH2

With limited information on the potential function of *Pf*AMC1, we used the AlphaFold 3.0 predicted structure as input for the 3D structure alignment tool Foldseek^37,55^. Even though no sequence homology was found through BLAST, Foldseek identified the OMM-localised human MTCH2 as a potential structural homologue of *Pf*AMC1. Interestingly, the potential homologues of *T. gondii* and *C. parvum* based on sequence analysis were not identified by Foldseek. In the case of *T. gondii* this is likely due to the regular large unstructured insertions while the unconventional groove is indeed not predicted for *C. parvum*. MTCH2 has been implicated in many different cellular processes ranging from lipid scrambling and homeostasis to apoptosis and OMM membrane protein insertion^56–58,73,74^. The molecular mechanism of lipid homeostasis of MTCH2 remains to be elucidated but its lipid scrambling function has been experimentally shown and computationally modelled^56,57,73,75^.

Interestingly, two independent groups showed that the deletion of *MTCH2* causes mitochondria to fragment and round up in mammalian cells, a phenotype that is comparable to 3HA tagging of *Pf*AMC1 (**Fig. 6**)^60,61,76^. Their data show that MTCH2 promotes mitochondrial fusion and depends on lysophosphatidic acid generated by ER- and OMM-resident glycerol-phosphate acyl transferase^61,76^. General consensus is that mitochondrial fusion does not occur in *Plasmodium*, mainly because of the predominantly single mitochondrion as well as the absence of identifiable mitofusins in the genome^77,78^. However, in a study from 2005 using live fluorescence imaging of asexual blood-stage *P. falciparum* potential mitochondrial fusion events were reported^79^. Moreover, we described the unexpected presence of multiple mitochondria *in P. falciparum* gametocytes^39,48^. These multiple mitochondria appear to be maintained during ookinete development in *P. berghei*^80^. It remains unclear what happens during oocyst development but the resulting sporozoites appear to have a single mitochondrion once again. While this could hint at mitochondrial fusion having a role during life-cycle progression, it would seem more likely that sporozoites simply include a single mitochondrion during their formation. Other studies have shown MTCH2 to be involved in apoptosis by binding the truncated apoptosis initiator Bid (tBid) and cascading Bak and Bax induced OMM pore formation, leading to cytochrome *c* release^74,81,82^. Intriguingly, the existence of programmed cell death in the form of apoptosis and autophagy in *Plasmodium* is again controversial and has not previously been studied during gametogenesis^83^. Furthermore, *Plasmodium* lacks obvious homologues of the proteins inducing the mitochondrial apoptotic pathway through MTCH2 (Bid, Bak, and Bax), although two putative Bax inhibitors have been identified (PF3D7_1248500 C PF3D7_1459800)^65^.

Lastly, MTCH2 was shown to be an α-helical and tail anchored OMM protein insertase. Mammalian cells also have the <u>m</u>itochondrial <u>im</u>port (MIM) complex, which is also shown to insert α-helical proteins into the OMM but this complex is not conserved in *Plasmodium*^84^.

Therefore, it remains to be elucidated how *Plasmodium* α-helical proteins are inserted into the OMM. It should be noted, however, that few α-helical OMM proteins are identified in *Plasmodium* and some canonical ones, such as OMP25, seem to be absent (BLAST). The only certain α-helical OMM to our knowledge is Fission 1 (PF3D7_1325600). The exact molecular mechanism of how MTCH2 exerts these diverse functions remains unknown. The predicted structural homology as well as the overlapping phenotype between *Pf*AMC1 and MTCH2 make it intriguing to speculate on a role of *Pf*AMC1 in either of these processes and provide a clear incentive for further research.

## Material and methods

### Bioinformatics

Protein structures were gathered from the PDB (https://www.rcsb.org/). The structural predictions of the respective proteins were obtained through the AlphaFold 3 Server^37^. These were then uploaded to the Foldseek webserver, searched against all available databases in 3DI/AA mode, to identify structural homologues^85^. The structural alignments were performed in Pymol 3.1 (Schrödinger) using the cealign command, which is very robust for proteins with very low sequence similarity. Electrostatic overviews were also generated in Pymol. Lastly, BLAST searches were performed in the NCBI webserver (https://blast.ncbi.nlm.nih.gov/Blast.cgi) using default parameters.

### *P. falciparum* culture, transfection and genotyping

All *P. falciparum* ABS cultures, WT (NF54), MitoRed, A1.1, and A1.2, were cultured in complete parasite culture media (RPMI1640 supplemented with 25 mM HEPES, 10% human type A serum, and 25 mM NaHCO_3_), where appropriate complemented with the indicated concentrations of glucosamine (glcN, Sigma, 1514)^39^. Parasites were maintained at 5% haematocrit (type O, Sanquin, the Netherlands) at 37 °C in low oxygen gas mixture (3% O_2_, 4% CO_2_). 60 μg of both Cas9-guide and linearized repair plasmids were transfected in ring-stage parasites^86,87^. In short, the precipitated Cas9-guide and repair plasmid were dissolved in 5 μL TE and mixed with cytomix (0.895% KCl, 0.0017% CaCl2, 0.076% EGTA, 0.102% MgCl2, 0.0871% K2HPO4, 0.068% KH2PO4, 0.708% HEPES) to a final volume of 30 µL and mixed with 0.3 mL packed ring-stage infected RBCs (iRBC). This mixture was electroporated (310 V, 950 μF) and allowed to recover for five hours before applying 2.5 nM WR99210 drug selection for five days. To exclude WT parasites from the population, 150 A1.1 parasites were sorted in the red channel (excitation 5610, detection filter 586/15 on the BD FACSAria (BD Biosciences), as line A1.1 harbours the red fluorescent *mito-mScar* cassette. To confirm absence of WT parasites in line A1.2, parasites were cultured on 2.5 mM glcN for 7 days followed by a diagnostic PCR (**Table S1**). Integration of both lines was confirmed by diagnostic PCR (**Table S1**).

### Plasmid construction

To generate the base vector that allows for 3xHA-glmS tagging with and without integrated HSP70-3 prom + mitochondrial targeting sequence + mScarlet (mito-mScar) (pRF138/pRF139), the 3xHA was amplified from pRF079 (**Table S1**) and cloned into pRF077/pRF079 using KpnI and NotI. The repair plasmid for line A1.1 was then generated by amplifying the 5’ and 3’ homology regions (HR) from genomic WT (NF54) DNA (**Table S1**) and cloned into pRF139 using XhoI/BamHI and EcoRI/NgoMIV. The Cas9-guide plasmids for line A1.1 were generated by annealing two pairs of oligonucleotides (**Table S1**) and cloning these into the pMLB626 plasmid (a kind gift from Marcus Lee). The base vector that allows for 3xHA-glmS tagging with selection on integration of DHFR (pRF289) was generated by amplifying the DHFR expression cassette from the pMLB626 plasmid (**Table S1**) Cas9-guide plasmid and cloning it into EcoRV cleaved pRF138. The PCR amplicon was cleaved with EcoRI and treated with DNA Polymerase I (Klenow, NEB, M0210) to create a blunt end. The repair plasmid for line A1.2 was generated by amplifying the 5’ and 3’ homology region (HR) from genomic WT (NF54) DNA (**Table S1**) and cloned into pRF289 using XhoI/BamHI and EcoRI/NgoMIV. The Cas9-guide plasmid for line A1.2 was generated by annealing oligonucleotides (**Table S1**) and cloning this into the pMLB626-DHODH plasmid (a kind gift from Marcus Lee).

### Growth assay

To obtain five-hour synchronous parasites, all lines were synchronised by loading a 50% haematocrit iRBC mixture containing segmenting schizonts on a 63% Percoll (GE Heakthcare, 17-0891-01) in PBS gradient and spinning at 1200*G for 20 minutes at 37°C. The intermediate layer was obtained and allowed to invade fresh RBCs for five hours. Ring-stage parasites were then selected by synchronising the culture with 5% sorbitol for 10 minutes at 37°C. The parasitaemia was then counted and independent cultures were set-up at 0.1-0.4%, with or without varying concentrations of glcN as required. Parasitaemia was monitored over seven days by fixing 10 µL of culture in 100 µL 0.25% glutaraldehyde (Panreac, A0589) in PBS. If blood smears indicated a parasitaemia of ∼5%, cultures were cut back to 0.3%. Parasites were left in fixation media for up to two weeks. On the day of readout, fixative was taken off and parasite pellet was resuspended in 1x SYBR-Green (Thermo Fisher, S7563). Samples were analysed on a Cytoflex flow cytometer (Beckman Coulter Cytoflex) using the 488 nm laser.

### SDS-PAGE western blot

Parasite cultures were harvested at indicated time points by resuspending packed iRBCs in 10x pellet volume 0.06% saponin (Sigma-Aldrich, S7900) in PBS for 10 minutes at 4°C. Saponin pellets were then resuspended in 2.5x pellet volume LDS sample buffer (GenScript, M00676) and 6.5x pellet volume Milli-Q (MQ), with 5% β-mercaptoethanol. Samples were heated at 70°C for 10 minutes, vortexed and centrifuged at 18.000*g for 3 minutes before loading on pre-cast Bis-Tris gels with varying percentages (GenScript Sure-PAGE^TM^ series). Proteins were transferred to a methanol activated PVDF membrane (BIO-RAD, 1620177) in transfer buffer (20% methanol with 10% Tris-Glycine (10x)) using a semi-dry Trans-Blot Turbo (Bio-Rad) at 25V for 30 minutes. The membrane was blocked in 2.5% milk powder in PBS for one hour at RT. Primary and secondary antibodies (**Table S2**) were diluted in 2.5% milk powder PBS and incubated overnight at 4°C (primaries) and 1 hour at RT (secondaries). Antibodies were washed four times five minutes. in PBS with 0.05% Tween20 (Sigma-Aldrich, P9416). The blots were imaged on a LAS4000 (ImageQuant) using Clarity^TM^ and Clarity Max^TM^ (BIO-RAD). Western blot densitometry was performed on ImageJ^88^.

### Drug sensitivity assay

The sensitivity of glcN treated and untreated WT and A1.1 parasites to several mitochondrial and non-mitochondrial targeting antimalarial compounds (compounds were a generous gift from TropIQ Health Sciences, Nijmegen) was determined using a replication assay as described in Schalkwijk *et al.*^89^. In short, 30 µL mixed ABS parasites were set up at 0.83% parasitaemia in 3% haematocrit in a 384-well plate. The compounds were serially diluted in dimethyl sulfoxide (DMSO) and RPMI1640 media to a final DMSO concentration of 0.1%. 30 µL of the serial diluted compounds was added to the parasites and incubated at 37°C for 72 hours in low oxygen gas mixture (3% O_2_, 4% CO_2_). Parasites were then lysed and stained for DNA in a SYBR-green lysis buffer (1:15,000 SYBR Green reagent (Life Technologies), 13.3 mM Tris-HCl, 3.3 mM EDTA, 0.067% TritonX-100, 0.0053% saponin). fluorescence intensity was quantified using BioTek Synergy 2 plate reader. Relative parasite growth was determined by comparing compound dilutions to DMSO (100% growth) and 1 µM dihydroartemisinin (DHA, 0% growth). GraphPad was used to determine the growth response curves with a variable slope model.

### Gametocyte induction and activation

Gametocytogenesis for all lines was similarly induced by culturing 1% trophozoites in Albumax supplemented media (RPMI1640, 2.5% AlbuMAXII^TM^ (Thermo Fisher, 11021-037), 25 mM NaHCO_3_) for 36 hours, before swapping back to complete parasite culturing media^90^. Gametocytes were cultured in a semi-automatic culturing system with twice a day media changes up until maturation at 13 days post induction^91^. All experiments and handling with gametocytes were performed with equipment preheated at 37°C. Exflagellation was triggered by mixing 10 µL activation media (50 μM xanthurenic acid (XA), Sigma-Aldrich, D120804) in complete parasite culture media) with 10 μL of mature stage V gametocyte cultures. This was loaded on a haemocytometer and incubated for 15 minutes at RT to quantify exflagellation events. Alternatively, it was loaded on a poly-L-lysine coated coverslip (Corning, 354085) for immunofluorescent microscopy (described below). To harvest activated samples for SDS-PAGE western blot analysis, packed iRBC were resuspended in 2.5x activation media and incubated for 20 minutes at RT, before being lysed in 0.06% saponin as described above.

### Immunofluorescent microscopy and analysis

When indicated, samples were stained with MitoTracker^TM^ by incubating 50 µL parasite culture with 450 µL MitoTracker^TM^ medium (100 nM MitoTracker^TM^ Orange CMTMRos (Thermo Fisher, M7510) in complete parasite culture media) for 30 minutes at 37°C. The samples were then washed and resuspended in 500 µL pre-heated complete parasite culture media and processed as the other samples. For all samples, a total of 20 µL 0.5% haematocrit iRBCs were loaded on a poly-L-lysine coated coverslip (Corning, 354085) and allowed to settle for 15 minutes at 37°C, before being fixed in 4% paraformaldehyde (Thermo Fisher, 28906) and 0.0075% glutaraldehyde (Panreac, #A0589,0010) in PBS for 20 minutes^92^. The coverslips were then washed in 1x PBS, permeabilised with 0.1% Triton X-100 in PBS for 10 minutes and blocked in 3% BSA for 1h at RT. They were then overnight incubated with primary antibodies (**Table S2**) in 3% BSA at 4°C, washed three times 5 minutes. and incubated with secondary antibodies (**Table S2**) for 1 hour at RT. The coverslips were then stained with 300 nM DAPI (Thermo Fisher, 62248) for 1 hour before being mounted onto microscope slides with VECTASHIELD (VWR, H-1000) and sealed with nail polish. Images were acquired on an LSM900 or 880 Airyscan confocal microscope (ZEISS) using the 63x oil objective. Laser power and detector sensibility were equal within experiments. Images were Airyscan processed before analysis using FIJI software.^88^ The brightness and contrast were individually adjusted for colocalisation purposes and histogram settings are depicted to allow fair comparison. The average HA bit value was determined by the sum of the total signal intensity (RawIntDen) within the cell body (determined by DIC cell perimeter) for each slice with a max intensity >1000 bit, divided by the number of slices with a max intensity >1000 bit. Arivis 4D Vision software was used for the 3D visualisation. Mitochondria of activated gametocytes were quantified as previously described.^93^ Briefly, using the Fiji plug-in MitochondriaAnalyser, 3D image stacks were pre-processed with the following parameters: subtract background (radius = 1.25 μm); sigma filter plus (radius = 2.0 μm); enhance local contrast (slope = 1.40 μm); and gamma correction (value = 0.9 μm)^53,88^. For thresholding, block size was set to 0.85 μm with a C value of 7. The functions despeckle and remove outliers with radius = 3 pixels was used for post-processing of thresholded images. Statistical significance was determined using one-way ANOVA test with Sidàk correction.

### Expansion microscopy

The samples were stained with MitoTracker^TM^ as described above. The parasites were fixed on poly-L-lysine coated coverslips as described above and then prepared for expansion microscopy as previously described^94^. In short, coverslips were incubated in AA/FA buffer (0.5% acrylamide (AA, Sigma, A4058), 0.38% formaldehyde (FA, Sigma, F8775) in PBS) for 5h at 37°C. Right before removing the coverslips from the AA/FA solution, a gelling solution was prepared by adding 0.5% tetramethylethylenediamine (TMED, Sigma, T7024) and 0.5% Ammonium persulfate (APS Sigma, A3678) to the U-ExM monome solution (MS, 23% w/v sodium acrylate (AK Scientific, R624), 10% w/v AA, N,N’-methylenbisacrylamide (BIS, Sigma, M1533) 0.1% w/v in PBS). This was then added on top of the coverslip and incubated at 37°C for 1 hour. The gels were then incubated with denaturation buffer (200 mM SDS, 200 mM NaCl, 50 mM Tris in MQ, pH 9) until they were completely detachment from the coverslips and then transferred into a 1.5 mL Eppendorf tube filled with fresh denaturation buffer and incubated for 90 minutes at 95°C. The gels were expanded overnight in MQ at RT and incubated with primary antibodies (**Table S2**) for 3h at 37°C. They were then washed three times in 0.06% PBS Tween-20 for 10 minutes and incubated with secondary antibodies and 600 nM DAPI (Thermo Fisher, 62248) for 2.5h at 37°C. The gels were cut into ∼5mm pieces and positioned on poly-D lysine (Gibco, A38904-01) coated coverslips. Images were acquired and analysed as described above.

## Acknowledgments

We thank all members of the Molecular & Cellular Parasitology team for the insightful discussions. We thank the Radboud Technology Center Microscopy and Radboud Technology Center Flow cytometry for the use of their facilities. We are grateful to Micheal Delves for sharing the anti-PF3D7_1325200 antibody, Marcus Lee for providing the pMLB626 CRISPR/Cas9 guide plasmid, and Koen Dechering and Tonnie Huijs for their support with the drug sensitivity assays. We thank Julie Verhoef for her assistance in the use of Arivis 4D Vision software and the generation of the 3xHA-glmS tagging base plasmids.

This work was supported by funding provided the Radboud University Medical Center (RIMLS018-009b to CB and TWAK), a short-term fellowship of the European Molecular Biology Organisation (SEG9997 to CB) and the Dutch Research Council (NWO, grant number: OCENW.M.21.087 to STL and TWAK). Ultra-expansion microscopy was funded by a Radboud Community of Infectious Disease grant (RCI Grant 2024 to STL and TWAK). ACK and ERSK were supported by core funding from the UKRI Medical Research Council (MC_UU_00028/2).

## Supplemental data

**Supplementary table 1.**
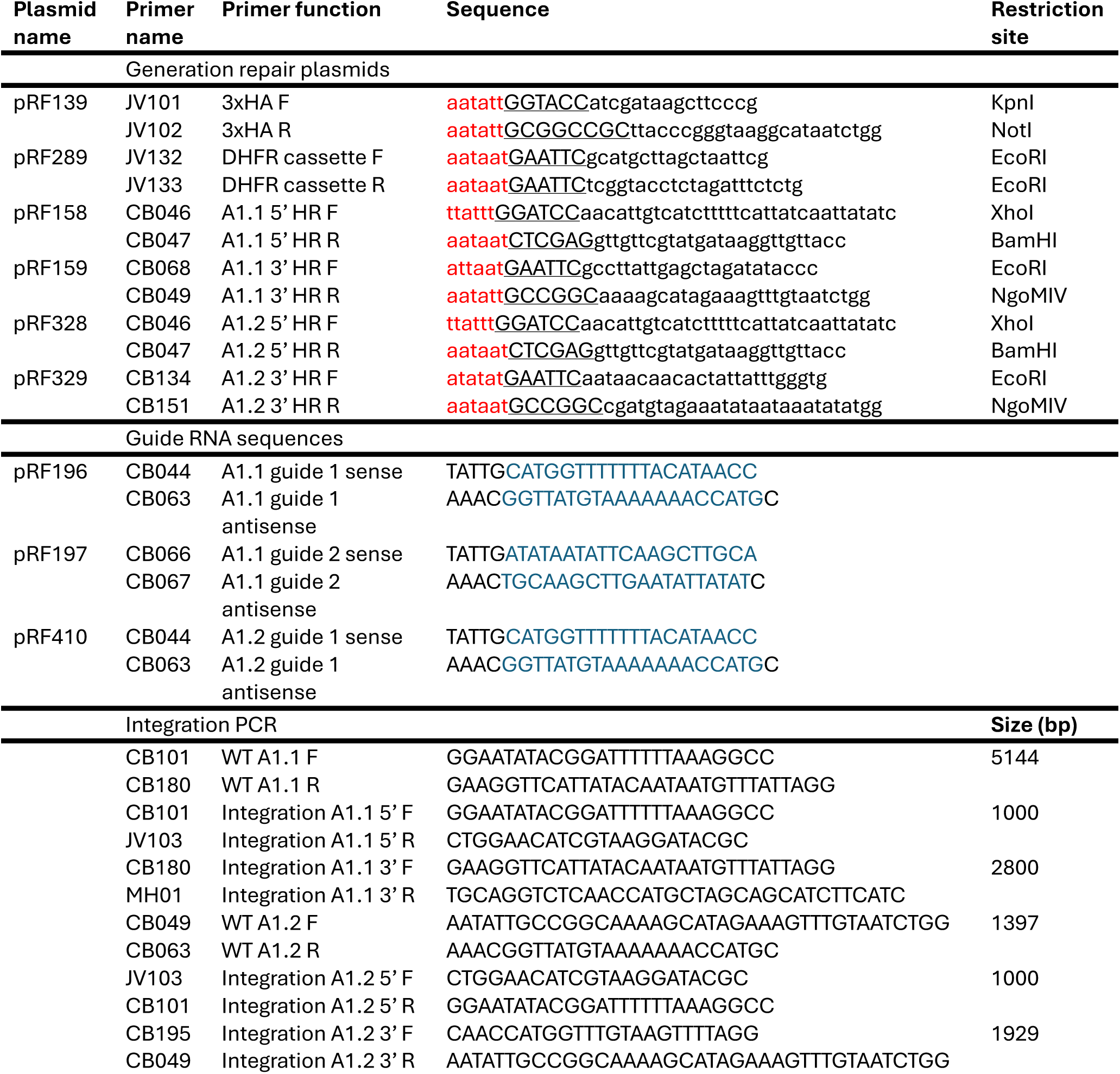
Primer and guide sequences used to generate repair and guide plasmids of lines A1.1 and A1.2.

**Supplementary table 2.** List of primary and secondary antibodies used for IFA/U-ExM and western blot.

| Antibody | Species | IFA/U-ExM dilution | WB dilution | Source |
| --- | --- | --- | --- | --- |
| Anti-HA IgG | Rat | 1:500/250 | 1:1000 | 11867423001 Roche (Merck) |
| Anti-Alpha Tubulin IgG | Mouse | 1:500/250 | - | MA1-19162, Invitrogen |
| Anti-PF3D7_1325200 (male marker) polyclonal | Rat | 1:10,000 | - | Kind gift from Dr. Michael Delves, LSHTM |
| Anti-HSP70 polyclonal | Rabbit | - | 1:2000 | SPC-186D, StressMarq Biosciences |
| Anti-Mouse IgG 700 IRDye Fluorophore | Goat | - | 1:5000 | 926-68070 IRDy, LI-COR |
| Anti-Rabbit IgG 680 IRDye Fluorophore | Goat | - | 1:5000 | 926-68071 IRDy, LI-COR |
| Anti-Rat IgG 800 IRDye Fluorophore | Goat | - | 1:5000 | 926-32219 IRDy, LI-COR |
| Anti-Rat IgG HRP | Mouse | - | 1:2000 | MCA189P, SEROTEC |
| Anti-rabbit IgG HRP | Goat | - | 1:2000 | P044801-2, Dako |
| Anti-Rat 488 IgG Alexa Fluorophore | Chicken | 1:500 | - | A-21470, Invitrogen |
| Anti- Rabbit IgG 488 Alexa Fluorophore | Chicken | 1:500 | - | A-21441 Invitrogen |
| Anti-Mouse IgG 647 Alexa Fluorophore | Donkey | 1:500 | - | A-31571, Invitrogen |
| Anti-Rat IgG 647 Alexa Fluorophore | Chicken | 1:500 | - | A-21472, Invitrogen |

**Supplementary table 3.** Percentage identity value of sequence alignment using UniProt Align.

|  | hAAC1 | PfAAC | Pf. AMC1 | Pb. AMC1 | Pv. AMC1 | Cp. AMC1 | Tg. AMC1 |
| --- | --- | --- | --- | --- | --- | --- | --- |
| H. AAC1 |  |  |  |  |  |  |  |
| Pf. AAC | 59.8 |  |  |  |  |  |  |
| Pf. AMC1 | 10.47 | 11.92 |  |  |  |  |  |
| Pb. AMC1 | 11.73 | 13.13 | 72.43 |  |  |  |  |
| Pv. AMC1 | 12.44 | 12.38 | 74.77 | 69.37 |  |  |  |
| Cp. AMC1 | 15.69 | 15.29 | 27 | 23.97 | 24.07 |  |  |
| Tg. AMC1 | 18.18 | 19.1 | 17.03 | 16.87 | 18.95 | 18.47 |  |

**Supplementary figure 1.**
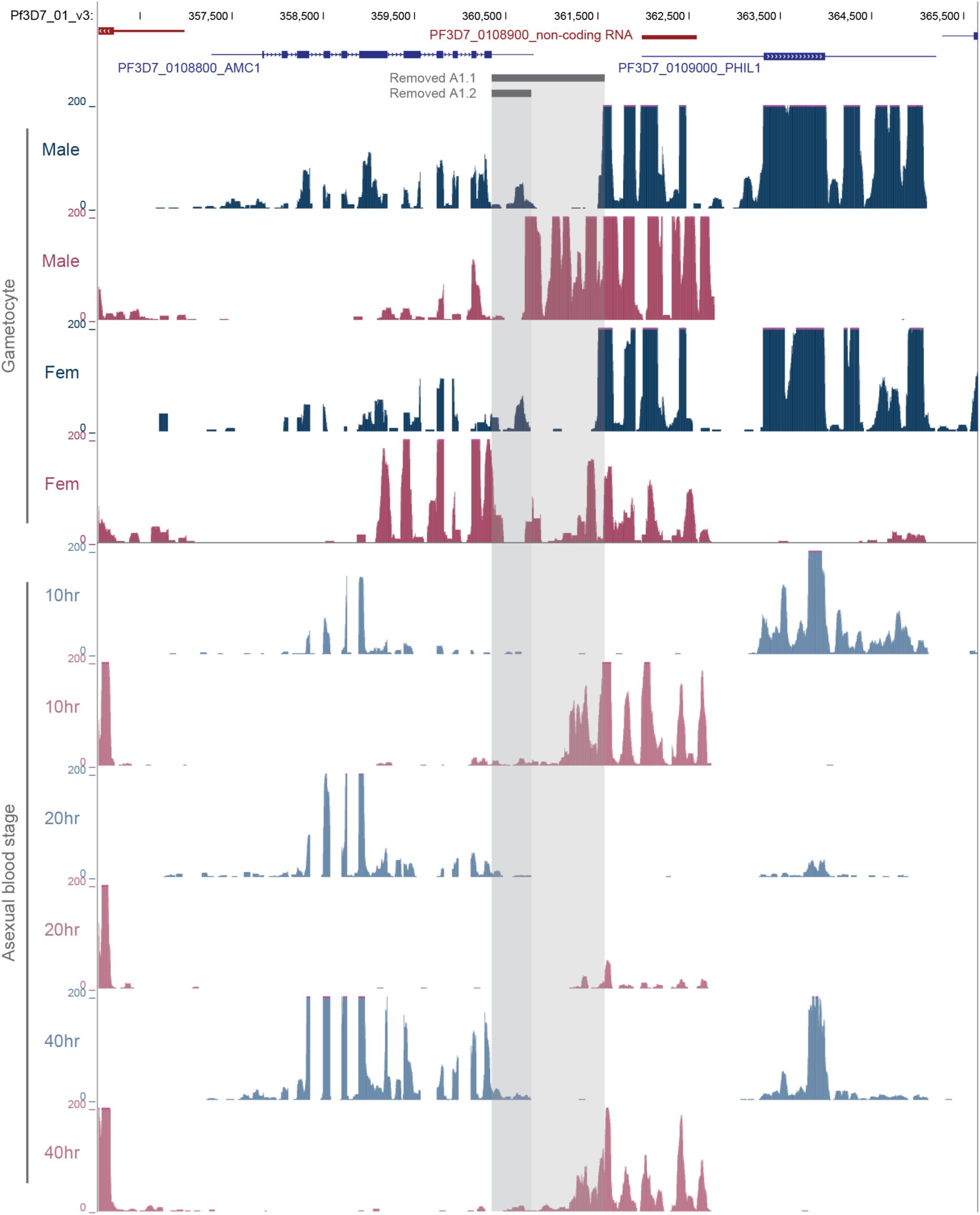
Expression profiles around the PfAMC1 locus in stage V gametocytes and ABS parasites. Profiles depict unique transcripts of the positive (blue) and negative (red) strand. The linear Y-axis is capped at 200 TPM to visualize PfAMC1 profiles accurately. The displayed gametocytes profiles are the unique reads from Lasonder et al. 201C^CC^. The displayed ABS profiles are the unique reads from Siegel et al. 2014^S^^5^.

**Supplementary figure 2.**
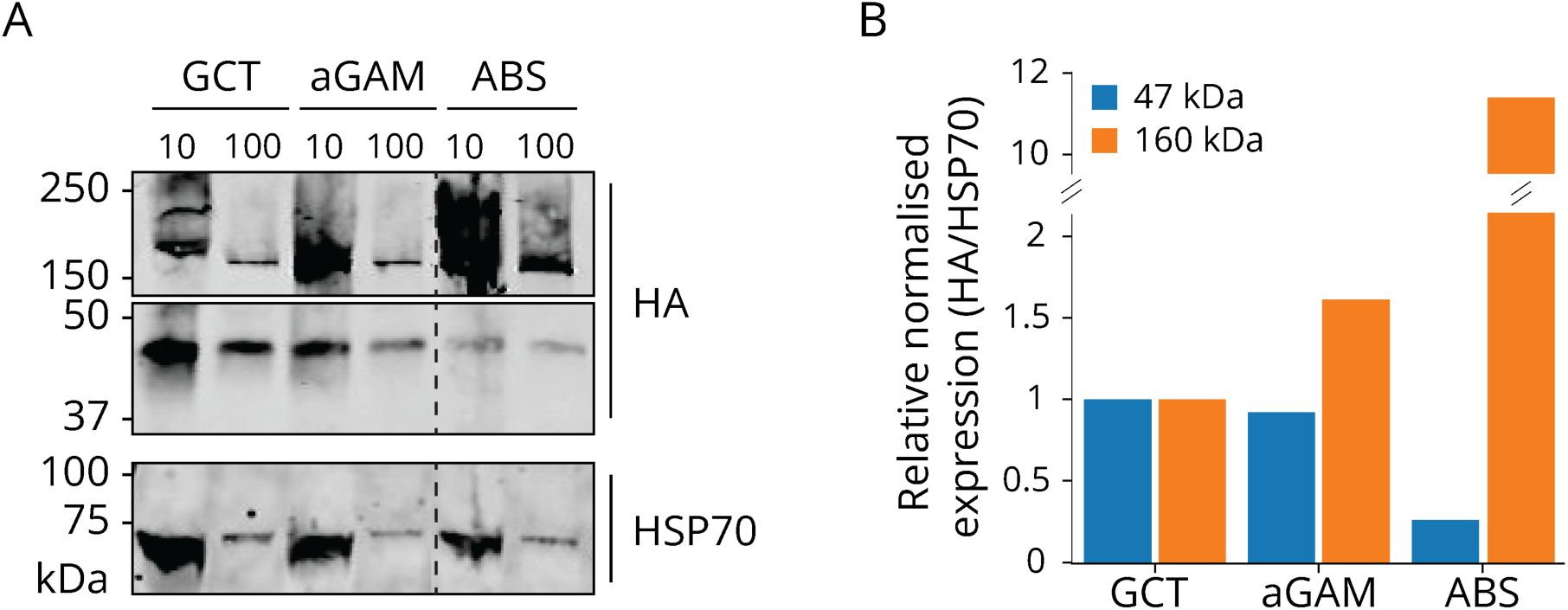
A) Western blot analysis of line A1.2 saponin lysed stage V gametocyte (GCT), 20 minutes XA activated gamete sample (aGAM), and mixed ABS stained with anti-HA and anti-HSP70. Samples were diluted in loading buffer at 1:10 as normal or 1:100 to more accurately determine the higher molecular weight band. B) The 47 and 160 kDa bands from the 1:100 dilution were normalised to their respective HSP70 bands and plotted to their respective intensity in stage V gametocytes.

**Supplementary figure 3.**
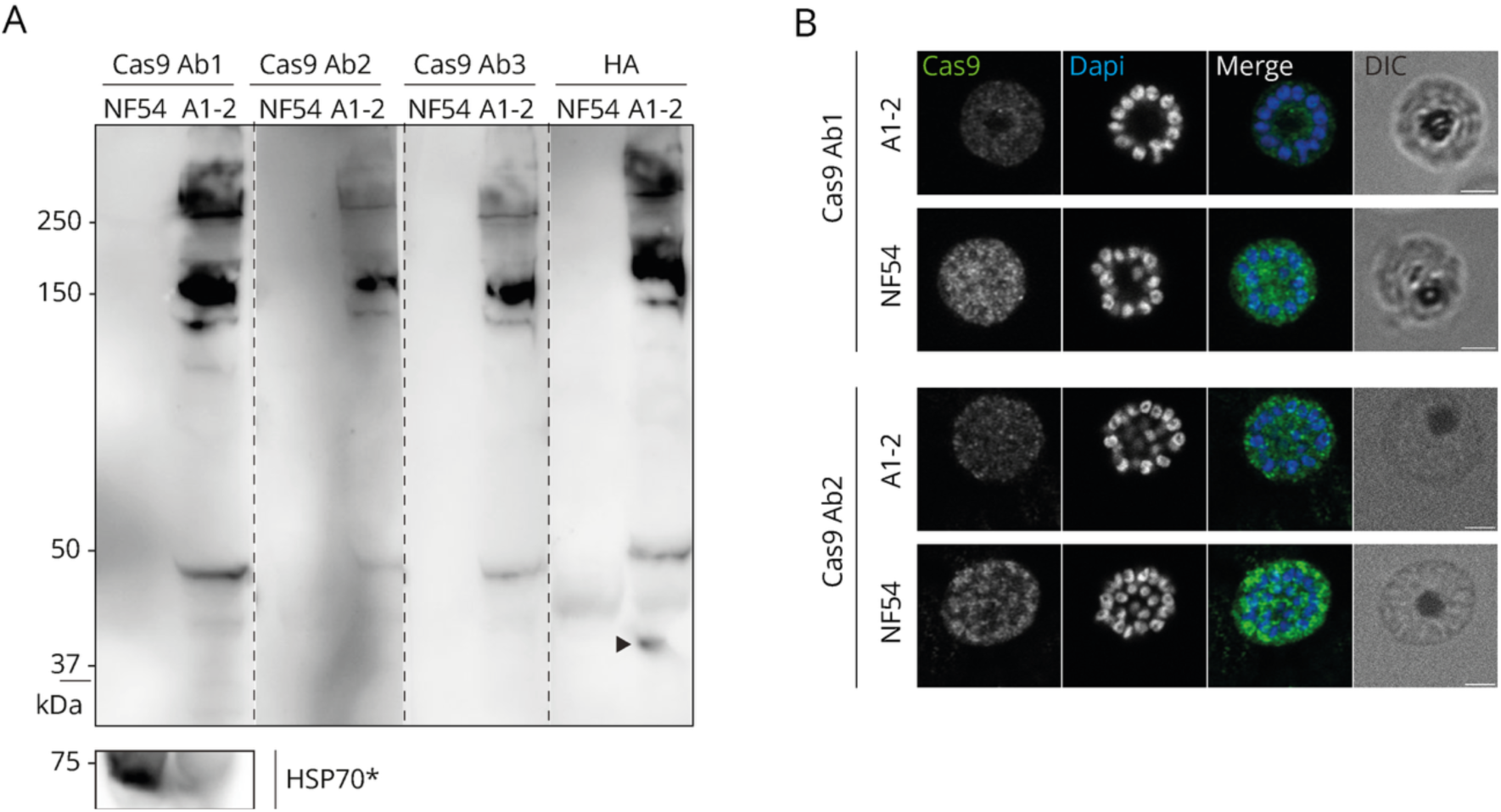
A1.2 has Cas3 expression. A) Western blot analysis of three different CasS and HSP70 antibodies on A1.2 mixed ABS. One lane was stained with the anti-HA antibodies. The black arrowhead indicates the additional AMC1 band, with an expected size of 45 kDa, absent in the CasS stained samples. *Same samples were loaded for each blot. B) Immunofluorescent images of two CasS antibodies on schizonts. The samples were stained with anti-CasS (green) antibodies and DAPI (blue). Scale bars, 2 µm.

**Supplementary figure 4.**
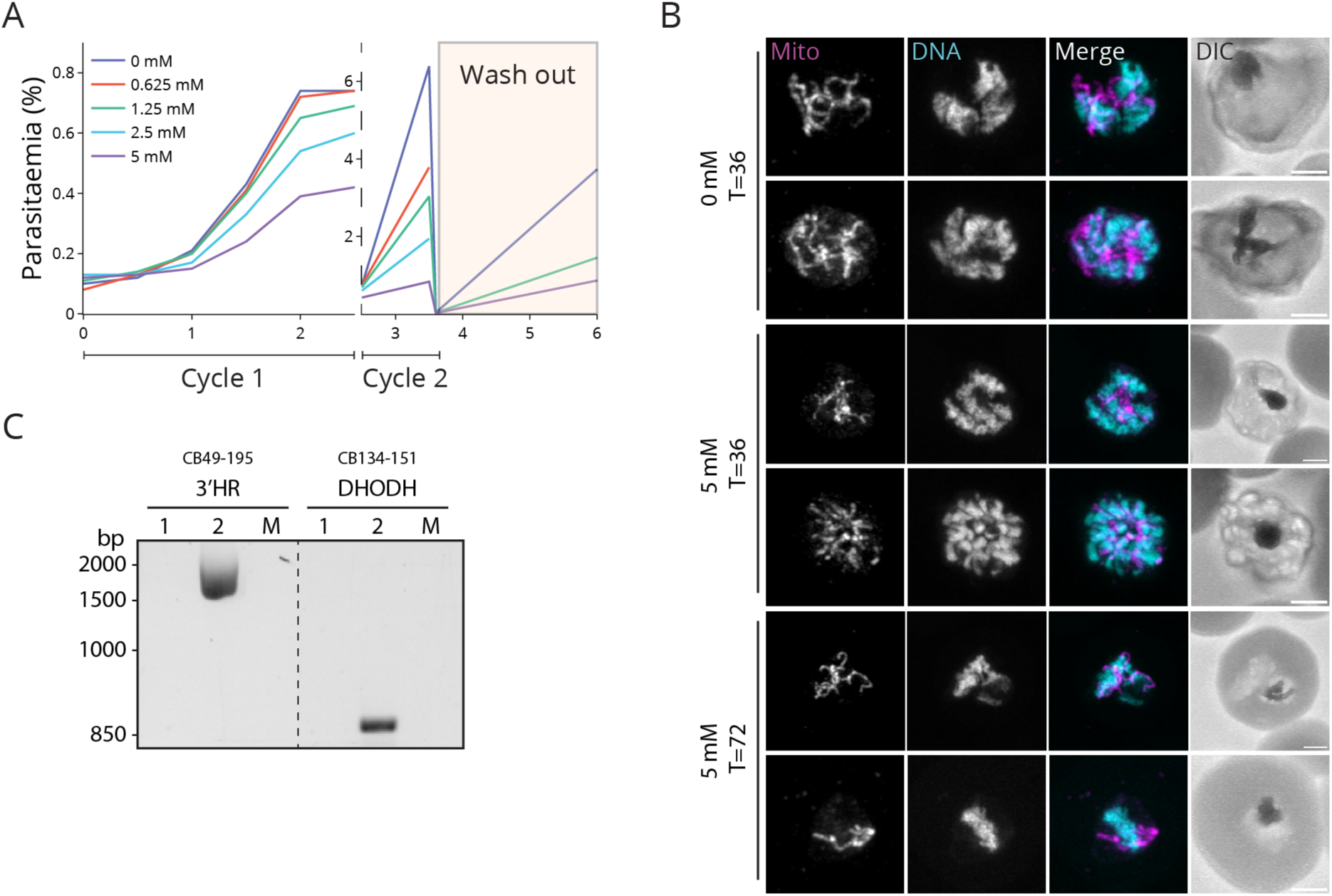
A) Titration growth assay comparing the growth rate of double sorbitol synchronised glcN treated A1.1 parasites, set-up at 0.1% parasitaemia. Cultures were cut back 1:50 after the second cycle and glcN media was replaced by complete parasite culture media (wash out). Samples were taken every 12 hours for the first cycle and then after another 24 and 48 hours. Parasitaemia was determined by flow cytometry, measuring DNA using SYBR-GreenTM. B) Immunofluorescent images of the growth assay cultures. The indicated samples were paraformaldehyde-glutaraldehyde fixed and stained with DAPI. Mitochondrial signal was obtained from the mito-mScar cassette present in line A1.1. C) Diagnostic PCR on gDNA from A1.1 (1), A1.2 (2), and MitoRed (M). Priming for 3’HR integration of line 2 (1S2S bp) and yDHODH (868 bp).

**Supplementary figure 5.**
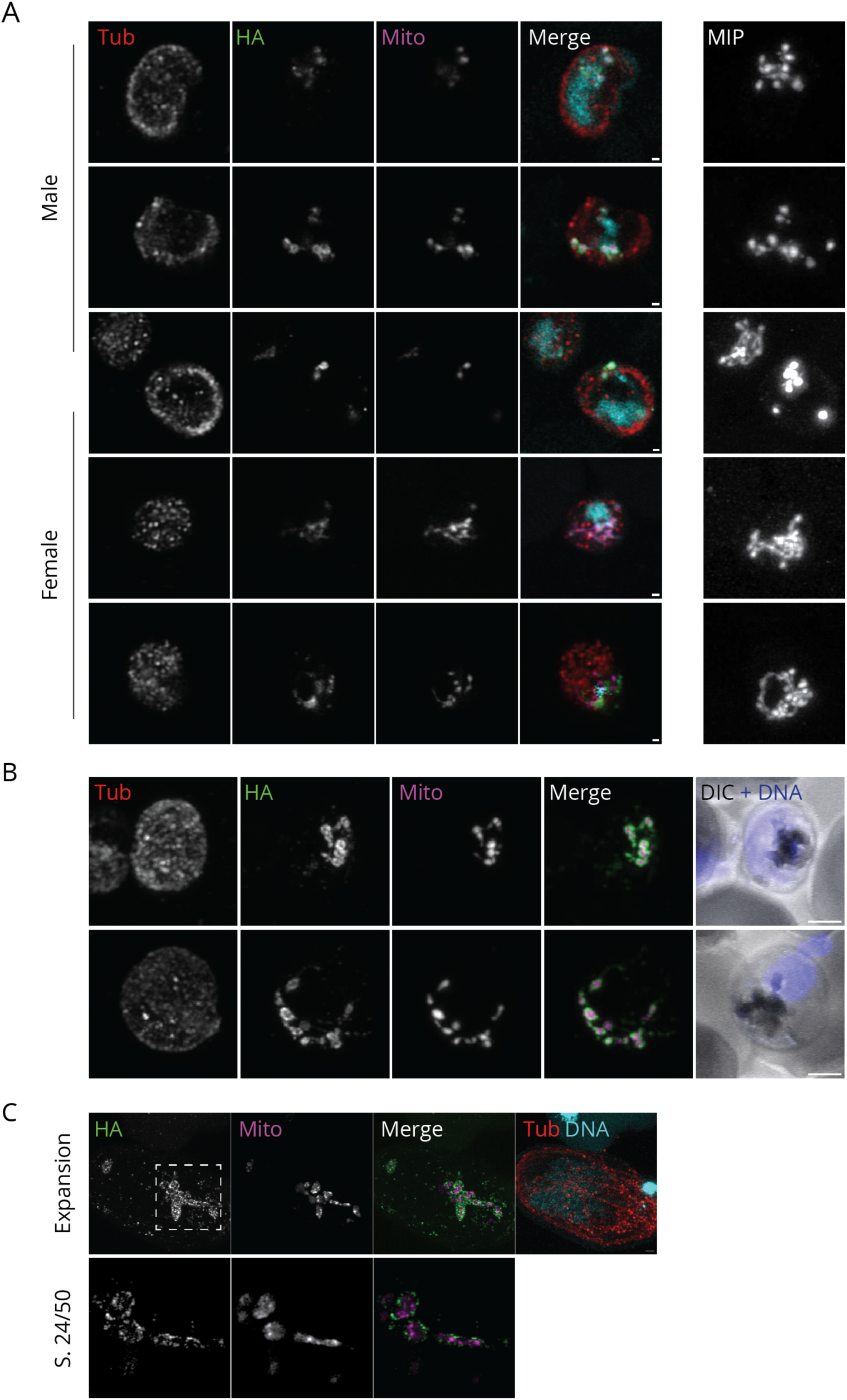
A&B) Immunofluorescent images of line A1.1 activated female (A) and male (A&B) gametocytes, stained with anti-HA (green), anti α-tubulin (red), and DAPI (DNA, cyan). The mitochondrial signal (mito, magenta) was captured from the remaining mito-mScar signal. Scale bars, 2 µm. Maximum intensity projections (MIP) of the complete Z-stack images are depicted to assess mitochondrial fragmentation. A) Individual slices of Z-stack images. B) Maximum intensity projection of Z-stack images. C) Expansion microscopy of A1.1 activated male gametocyte stained with anti-HA (green), anti α-tubulin (red), DAPI, and MitoTracker^TM^ Orange. Dashed square indicates single slice zoom-in.

**Supplementary figure 6.**
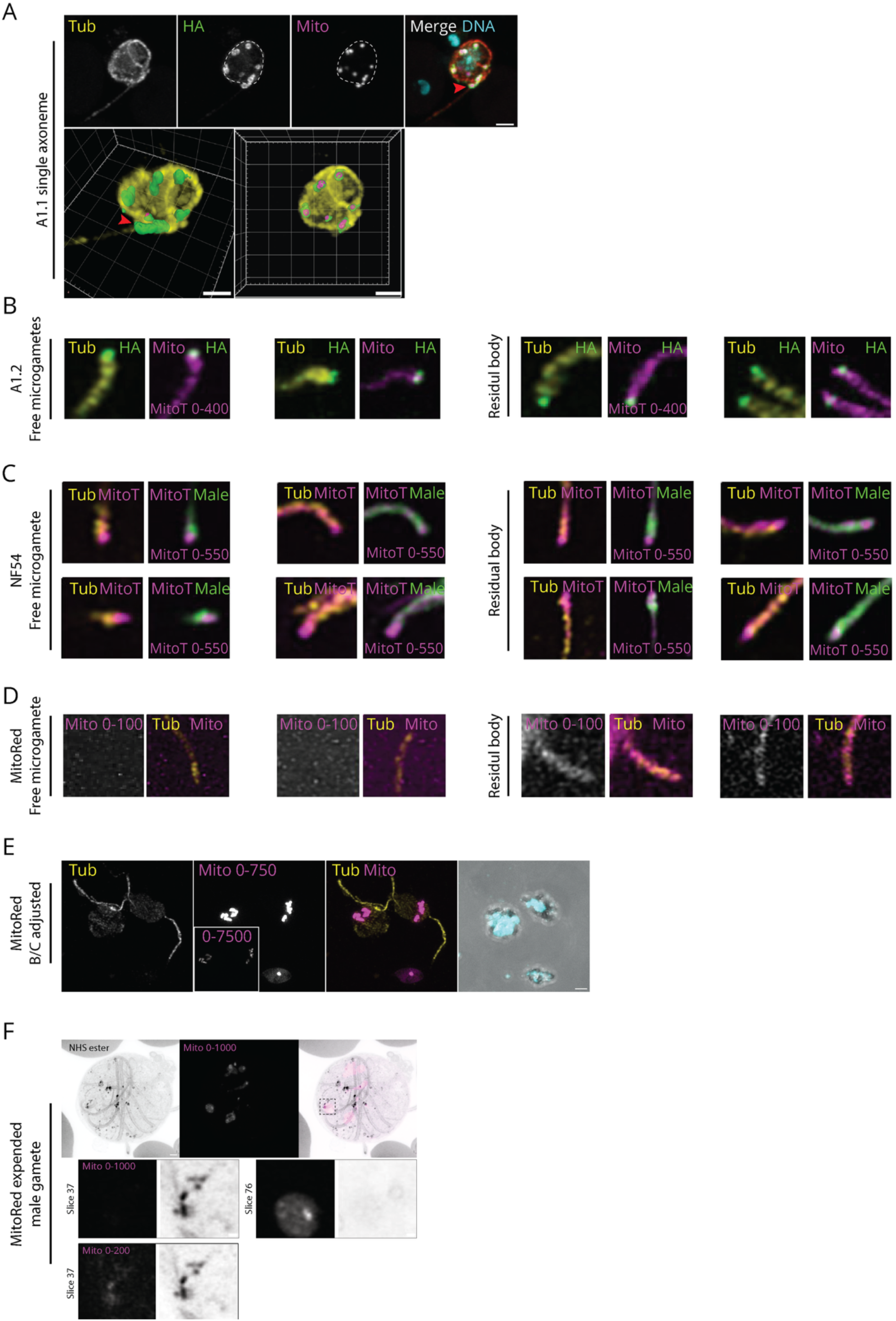
A) 3D visualisation of an A1.1 activated male gametocyte with a single extended axoneme. Cell periphery determined by α-tubulin signal is indicated with a white dashed line. Red arrowheads indicate a mitochondrial fragment in the same plane as the axoneme. Scale bars, 2 µm. B-F) Analysis of microgametes of different lines to assess the polarity and origin of the tubulin free compartment. Numbers in the MitoTracker^TM^ channel indicate histogram settings. B) A1.2 microgametes stained with anti-α-tubulin, anti-HA, MitoTracker^TM^ Orange, and DAPI. C) WT microgametes stained with anti-α-tubulin, anti-male marker (Male), MitoTracker^TM^ Orange, and DAPI. D) MitoRed microgametes stained with anti-α-tubulin and DAPI. Mitochondrial signal was captured from the mito-mScar signal. E) Brightness and contrast (B/C) adjusted MitoRed exflagellating male gamete to assess mito-mScar background signal compared to the MitoTracker^TM^ stained expansion microscopy (Fig. 5E and Supp. fig. 5F). F) Expanded MitoRed activated male gamete stained with MitoTracker^TM^ Orange and NHS ester. Zoom-ins from the dashed square indicate different slices to assess basal body and mitochondrial co-localisation.

**Supplementary figure 7.**
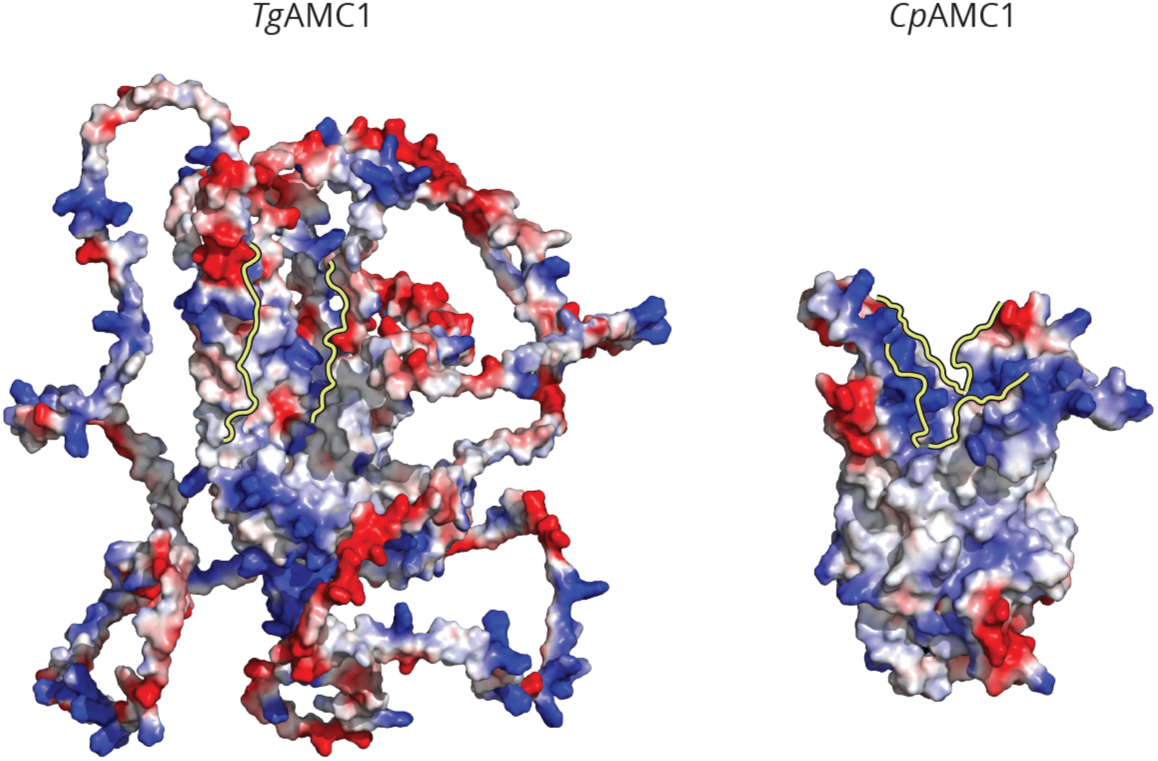
Electrostatic overview of the predicted T. gondii (Tg) and C. parvum (Cp) AMC1 structure with the groove highlighted by the yellow lines. The loop structures of T. gondii AMC1 have no defined fold.

## Supplementary video available on GitHub (CasBosh/AMC1)

**Supplementary video 1.** Corresponding to cell in figure 4G with DNA (cyan), tubulin (red), mitochondria (magenta), AMC1 (green).

**Supplementary video 2.** Corresponding to cell in supplemental figure 6A with DNA (cyan), tubulin (red), mitochondria (magenta), AMC1 (green).

## References

1. World Health Organization. World Malaria Report 2024. (2024).

2. Vaidya, A. B. & Mather, M. W. Mitochondrial evolution and functions in malaria parasites. Annu. Rev. Microbiol. 63, 249–267 (2009).

3. Evers, F. et al. Composition and stage dynamics of mitochondrial complexes in Plasmodium falciparum. Nat. Commun. 12, 3820 (2021).

4. Gujjar, R. et al. Identification of a metabolically stable triazolopyrimidine-based dihydroorotate dehydrogenase inhibitor with antimalarial activity in mice. J. Med. Chem. 52, 1864–1872 (2009).

5. Fry, M. & Pudney, M. Site of action of the antimalarial hydroxynaphthoquinone, 2-[trans-4- (4’-chlorophenyl) cyclohexyl]-3-hydroxy-1,4-naphthoquinone (566C80). Biochem. Pharmacol. 43, 1545–1553 (1992).

6. Srivastava, I. K., Rottenberg, H. & Vaidya, A. B. Atovaquone, a broad spectrum antiparasitic drug, collapses mitochondrial membrane potential in a malarial parasite. J. Biol. Chem. 272, 3961–3966 (1997).

7. Fry, M., Webb, E. & Pudney, M. Effect of mitochondrial inhibitors on adenosinetriphosphate levels in Plasmodium falciparum. Comp. Biochem. Physiol. B 96, 775–782 (1990).

8. Painter, H. J., Morrisey, J. M., Mather, M. W. & Vaidya, A. B. Specific role of mitochondrial electron transport in blood-stage Plasmodium falciparum. Nature 446, 88–91 (2007).

9. Sadik, M., Afsar, M., Ramachandran, R. & Habib, S. [Fe-S] biogenesis and unusual assembly of the ISC scaffold complex in the Plasmodium falciparum mitochondrion. Mol. Microbiol. 116, 606–623 (2021).

10. Tian, H.-F., Feng, J.-M. & Wen, J.-F. The evolution of cardiolipin biosynthesis and maturation pathways and its implications for the evolution of eukaryotes. BMC Evol. Biol. 12, 32 (2012).

11. Jenkins, B. J. et al. Characterization of a Plasmodium falciparum Orthologue of the Yeast Ubiquinone-Binding Protein, Coq10p. PLoS One 11, e0152197 (2016).

12. Ellenrieder, L. et al. Dual Role of Mitochondrial Porin in Metabolite Transport across the Outer Membrane and Protein Transfer to the Inner Membrane. Mol. Cell 73, 1056–1065.e7 (2019).

13. Kunji, E. R. S., King, M. S., Ruprecht, J. J. & Thangaratnarajah, C. The SLC25 Carrier Family: Important Transport Proteins in Mitochondrial Physiology and Pathology. Physiology (Bethesda). 35, 302–327 (2020).

14. Ruprecht, J. J. & Kunji, E. R. S. Structural Mechanism of Transport of Mitochondrial Carriers. Annu. Rev. Biochem. 90, 535–558 (2021).

15. Pebay-Peyroula, E. et al. Structure of mitochondrial ADP/ATP carrier in complex with carboxyatractyloside. Nature 426, 39–44 (2003).

16. Robinson, A. J., Overy, C. & Kunji, E. R. S. The mechanism of transport by mitochondrial carriers based on analysis of symmetry. Proceedings of the National Academy of Sciences 105, 17766–17771 (2008).

17. Robinson, A. J. & Kunji, E. R. S. Mitochondrial carriers in the cytoplasmic state have a common substrate binding site. Proc. Natl. Acad. Sci. U. S. A. 103, 2617–2622 (2006).

18. Kunji, E. R. S. & Robinson, A. J. The conserved substrate binding site of mitochondrial carriers. Biochim. Biophys. Acta Bioenerg. 1757, 1237–1248 (2006).

19. Ruprecht, J. J. et al. The Molecular Mechanism of Transport by the Mitochondrial ADP/ATP Carrier. Cell 176, 435–447.e15 (2019).

20. King, M. S., Kerr, M., Crichton, P. G., Springett, R. & Kunji, E. R. S. Formation of a cytoplasmic salt bridge network in the matrix state is a fundamental step in the transport mechanism of the mitochondrial ADP/ATP carrier. Biochim. Biophys. Acta Bioenerg. 1857, 14–22 (2016).

21. Ruprecht, J. J. et al. Structures of yeast mitochondrial ADP/ATP carriers support a domain-based alternating-access transport mechanism. Proceedings of the National Academy of Sciences 111, E426–E434 (2014).

22. van Dooren, G. G., Stimmler, L. M. & McFadden, G. I. Metabolic maps and functions of the Plasmodium mitochondrion. FEMS Microbiol. Rev. 30, 596–630 (2006).

23. Nozawa, A., Fujimoto, R., Matsuoka, H., Tsuboi, T. & Tozawa, Y. Cell-free synthesis, reconstitution, and characterization of a mitochondrial dicarboxylate-tricarboxylate carrier of Plasmodium falciparum. Biochem. Biophys. Res. Commun. 414, 612–617 (2011).

24. Nozawa, A. et al. Characterization of mitochondrial carrier proteins of malaria parasite Plasmodium falciparum based on in vitro translation and reconstitution. Parasitol. Int. 79, 102160 (2020).

25. Weiner, J. & Kooij, T. W. A. Phylogenetic profiles of all membrane transport proteins of the malaria parasite highlight new drug targets. Microb. Cell 3, 511–521 (2016).

26. Bushell, E. et al. Functional Profiling of a Plasmodium Genome Reveals an Abundance of Essential Genes. Cell 170, 260–272.e8 (2017).

27. Zhang, M. et al. Uncovering the essential genes of the human malaria parasite Plasmodium falciparum by saturation mutagenesis. Science 360, (2018).

28. van Strien, J. et al. Analysis of Complexome Profiles with the Gaussian Interaction Profiler (GIP) Reveals Novel Protein Complexes in Plasmodium falciparum. J. Proteome Res. 23, 4467–4479 (2024).

29. van Esveld, S. L. et al. A Prioritized and Validated Resource of Mitochondrial Proteins in Plasmodium Identifies Unique Biology. mSphere e0061421 (2021) doi:10.1128/mSphere.00614-21.

30. Kulawiak, B. et al. The mitochondrial protein import machinery has multiple connections to the respiratory chain. Biochim. Biophys. Acta 1827, 612–626 (2013).

31. Pierrel, F. et al. Coa1 links the Mss51 post-translational function to Cox1 cofactor insertion in cytochrome c oxidase assembly. EMBO J. 26, 4335–4346 (2007).

32. Mick, D. U. et al. Shy1 couples Cox1 translational regulation to cytochrome c oxidase assembly. EMBO J. 26, 4347–4358 (2007).

33. Aurrecoechea, C. et al. PlasmoDB: a functional genomic database for malaria parasites. Nucleic Acids Res. 37, D539–43 (2009).

34. Letunic, I., Khedkar, S. & Bork, P. SMART: recent updates, new developments and status in 2020. Nucleic Acids Res. 49, D458–D460 (2021).

35. Jones, P. et al. InterProScan 5: genome-scale protein function classification. Bioinformatics 30, 1236–1240 (2014).

36. Sichrovsky, M. et al. Molecular basis of pyruvate transport and inhibition of the human mitochondrial pyruvate carrier. Sci. Adv. 11, 1489 (2025).

37. Abramson, J. et al. Accurate structure prediction of biomolecular interactions with AlphaFold 3. Nature 630, 493–500 (2024).

38. Prommana, P. et al. Inducible knockdown of Plasmodium gene expression using the glmS ribozyme. PLoS One 8, e73783 (2013).

39. Verhoef, J. M. J. et al. Detailing organelle division and segregation in Plasmodium falciparum. J. Cell Biol. 223, (2024).

40. Ghorbal, M. et al. Genome editing in the human malaria parasite Plasmodium falciparum using the CRISPR-Cas9 system. Nat. Biotechnol. 32, 819–821 (2014).

41. Ganesan, S. M. et al. Yeast dihydroorotate dehydrogenase as a new selectable marker for Plasmodium falciparum transfection. Mol. Biochem. Parasitol. 177, 29–34 (2011).

42. Pietsch, E. et al. A patatin-like phospholipase is important for mitochondrial function in malaria parasites. mBio 14, e0171823 (2023).

43. Sheokand, P., Mühleip, A. & Sheiner, L. Plasmodium falciparum mitochondrial complex III, the target of atovaquone, is essential for progression to the transmissible sexual stages. bioRxiv 2024.01.09.574740 (2024) doi:10.1101/2024.01.09.574740.

44. Goodman, C. D., Buchanan, H. D. & McFadden, G. I. Is the Mitochondrion a Good Malaria Drug Target? Trends Parasitol. 33, 185–193 (2017).

45. Mounkoro, P., Michel, T. & Meunier, B. Revisiting the mode of action of the antimalarial proguanil using the yeast model. Biochem. Biophys. Res. Commun. 534, 94–98 (2021).

46. Lelièvre, J., Berry, A. & Benoit-Vical, F. Artemisinin and chloroquine: do mode of action and mechanism of resistance involve the same protagonists? Curr. Opin. Investig. Drugs 8, 117–124 (2007).

47. de Vries, L. E. et al. Preclinical characterization and target validation of the antimalarial pantothenamide MMV693183. Nat. Commun. 13, 2158 (2022).

48. Evers, F. et al. Comparative 3D ultrastructure of Plasmodium falciparum gametocytes. Nat. Commun. 16, 69 (2025).

49. Sinden, R. E., Canning, E. U., Bray, R. S. & Smalley, M. E. Gametocyte and gamete development in Plasmodium falciparum. Proc. R. Soc. Lond. B Biol. Sci. 201, 375–399 (1978).

50. Creasey, A. et al. Maternal inheritance of extrachromosomal DNA in malaria parasites. Mol. Biochem. Parasitol. 65, 95–98 (1994).

51. Sparkes, P. C. et al. Mitochondrial ATP synthesis is essential for efficient gametogenesis in Plasmodium falciparum. Commun. Biol. 7, 1525 (2024).

52. Rashpa, R. & Brochet, M. Expansion microscopy of Plasmodium gametocytes reveals the molecular architecture of a bipartite microtubule organisation centre coordinating mitosis with axoneme assembly. PLoS Pathog. 18, e1010223 (2022).

53. Chaudhry, A., Shi, R. & Luciani, D. S. A pipeline for multidimensional confocal analysis of mitochondrial morphology, function, and dynamics in pancreatic β-cells. Am. J. Physiol. Endocrinol. Metab. 318, E87–E101 (2020).

54. Jumper, J. et al. Highly accurate protein structure prediction with AlphaFold. Nature 596, 583–589 (2021).

55. Barrio-Hernandez, I. et al. Clustering predicted structures at the scale of the known protein universe. Nature 622, 637–645 (2023).

56. Li, D., Rocha-Roa, C., Schilling, M. A., Reinisch, K. M. & Vanni, S. Lipid scrambling is a general feature of protein insertases. bioRxiv: the preprint server for biology Preprint at 10.1101/2023.09.01.555937 (2023).

57. Bartoš, L., Menon, A. K. & Vácha, R. Insertases Scramble Lipids: Molecular Simulations of MTCH2. bioRxiv: the preprint server for biology Preprint at 10.1101/2023.08.14.553169 (2023).

58. Guna, A. et al. MTCH2 is a mitochondrial outer membrane protein insertase. Science 378, 317–322 (2022).

59. Grinberg, M. et al. Mitochondrial carrier homolog 2 is a target of tBID in cells signaled to die by tumor necrosis factor alpha. Mol. Cell. Biol. 25, 4579–4590 (2005).

60. Bahat, A. et al. MTCH2-mediated mitochondrial fusion drives exit from naïve pluripotency in embryonic stem cells. Nat. Commun. 9, 5132 (2018).

61. Labbé, K. et al. The modified mitochondrial outer membrane carrier MTCH2 links mitochondrial fusion to lipogenesis. J. Cell Biol. 220, (2021).

62. Palmieri, F., Pierri, C. L., De Grassi, A., Nunes-Nesi, A. & Fernie, A. R. Evolution, structure and function of mitochondrial carriers: a review with new insights. Plant J. 66, 161–181 (2011).

63. Palmieri, F. & Monné, M. Discoveries, metabolic roles and diseases of mitochondrial carriers: A review. Biochim. Biophys. Acta 1863, 2362–2378 (2016).

64. Ruprecht, J. J. & Kunji, E. R. S. The SLC25 Mitochondrial Carrier Family: Structure and Mechanism. Trends Biochem. Sci. 45, 244–258 (2020).

65. Amos, B. et al. VEuPathDB: the eukaryotic pathogen, vector and host bioinformatics resource center. Nucleic Acids Res. 50, D898–D911 (2022).

66. Lasonder, E. et al. Integrated transcriptomic and proteomic analyses of P. falciparum gametocytes: molecular insight into sex-specific processes and translational repression. Nucleic Acids Res. 44, 6087–6101 (2016).

67. Saini, E. et al. Photosensitized INA-Labelled protein 1 (PhIL1) is novel component of the inner membrane complex and is required for Plasmodium parasite development. Sci. Rep. 7, 15577 (2017).

68. Saini, E. et al. Plasmodium falciparum PhIL1-associated complex plays an essential role in merozoite reorientation and invasion of host erythrocytes. PLoS Pathog. 17, e1009750 (2021).

69. Parkyn Schneider, M. et al. Disrupting assembly of the inner membrane complex blocks Plasmodium falciparum sexual stage development. PLoS Pathog. 13, e1006659 (2017).

70. Francia, M. E., Dubremetz, J.-F. & Morrissette, N. S. Basal body structure and composition in the apicomplexans Toxoplasma and Plasmodium. Cilia 5, 3 (2015).

71. Hair, M. et al. Atypical flagella assembly and haploid genome coiling during male gamete formation in Plasmodium. Nat. Commun. 14, 8263 (2023).

72. Talman, A. M. et al. Proteomic analysis of the Plasmodium male gamete reveals the key role for glycolysis in flagellar motility. Malar. J. 13, 315 (2014).

73. Rottiers, V. et al. MTCH2 is a conserved regulator of lipid homeostasis. Obesity (Silver Spring). 25, 616–625 (2017).

74. Zaltsman, Y. et al. MTCH2/MIMP is a major facilitator of tBID recruitment to mitochondria. Nat. Cell Biol. 12, 553–562 (2010).

75. Chourasia, S. et al. MTCH2 controls energy demand and expenditure to fuel anabolism during adipogenesis. 10.1038/s44318-024-00335-7 doi:10.1038/s44318-024-00335-7.

76. Goldman, A. et al. MTCH2 cooperates with MFN2 and lysophosphatidic acid synthesis to sustain mitochondrial fusion. EMBO Rep. 25, 45–67 (2024).

77. Stanway, R. R. et al. Organelle segregation into Plasmodium liver stage merozoites. Cell. Microbiol. 13, 1768–1782 (2011).

78. Voleman, L. & Doležal, P. Mitochondrial dynamics in parasitic protists. PLoS Pathog. 15, e1008008 (2019).

79. van Dooren, G. G. et al. Development of the endoplasmic reticulum, mitochondrion and apicoplast during the asexual life cycle of Plasmodium falciparum. Mol. Microbiol. 57, 405–419 (2005).

80. Darif, N. et al. Cellular Hallmarks From Volume Electron Microscopy Reveal Developmental Progression of Plasmodium Ookinetes. Adv. Sci. (Weinh). 13, (2026).

81. Hung, C.-L., Chang, H.-H., Lee, S. W. & Chiang, Y.-W. Stepwise activation of the pro-apoptotic protein Bid at mitochondrial membranes. Cell Death Differ. 28, 1910–1925 (2021).

82. Shamas-Din, A. et al. Multiple partners can kiss-and-run: Bax transfers between multiple membranes and permeabilizes those primed by tBid. Cell Death Dis. 5, e1277 (2014).

83. Sena-Dos-Santos, C. et al. Unraveling Cell Death Pathways during Malaria Infection: What Do We Know So Far? Cells 10, (2021).

84. Eaglesfield, R. & Tokatlidis, K. Targeting and Insertion of Membrane Proteins in Mitochondria. Front. Cell Dev. Biol. 9, 803205 (2021).

85. van Kempen, M. et al. Fast and accurate protein structure search with Foldseek. Nat. Biotechnol. 42, 243–246 (2024).

86. Wu, Y., Sifri, C. D., Lei, H. H., Su, X. Z. & Wellems, T. E. Transfection of Plasmodium falciparum within human red blood cells. Proc. Natl. Acad. Sci. U. S. A. 92, 973–977 (1995).

87. Crabb, B. S. & Cowman, A. F. Characterization of promoters and stable transfection by homologous and nonhomologous recombination in Plasmodium falciparum. Proc. Natl. Acad. Sci. U. S. A. 93, 7289–7294 (1996).

88. Schindelin, J. et al. Fiji: an open-source platform for biological-image analysis. Nat. Methods 9, 676–682 (2012).

89. Schalkwijk, J. et al. Antimalarial pantothenamide metabolites target acetyl-coenzyme A biosynthesis in Plasmodium falciparum. Sci. Transl. Med. 11, (2019).

90. Graumans, W. et al. AlbuMAX supplemented media induces the formation of transmission-competent P. falciparum gametocytes. Mol. Biochem. Parasitol. 259, 111634 (2024).

91. Ponnudurai, T., Lensen, A. H., Meis, J. F. & Meuwissen, J. H. Synchronization of Plasmodium falciparum gametocytes using an automated suspension culture system. Parasitology 93 (Pt 2), 263–274 (1986).

92. Tonkin, C. J. et al. Localization of organellar proteins in Plasmodium falciparum using a novel set of transfection vectors and a new immunofluorescence fixation method. Mol. Biochem. Parasitol. 137, 13–21 (2004).

93. Tassan-Lugrezin, S. et al. Building without the architect: the dispensable role of MICOS in de novo cristae formation in malaria parasites. bioRxiv 2025.10.13.682069 (2025) doi:10.1101/2025.10.13.682069.

94. Gambarotto, D., Hamel, V. & Guichard, P. Ultrastructure expansion microscopy (U-ExM). Methods Cell Biol. 161, 57–81 (2021).

95. Siegel, T. N. et al. Strand-specific RNA-Seq reveals widespread and developmentally regulated transcription of natural antisense transcripts in Plasmodium falciparum. BMC Genomics 15, 150 (2014).

